# Keratins couple with the nuclear lamina and regulate proliferation in colonic epithelial cells

**DOI:** 10.1101/2020.06.22.164467

**Authors:** Carl-Gustaf A. Stenvall, Joel H. Nyström, Ciarán Butler-Hallissey, Stephen A. Adam, Roland Foisner, Karen M. Ridge, Robert D. Goldman, Diana M. Toivola

**Affiliations:** Cell Biology, Biosciences, Faculty of Science and Engineering, Åbo Akademi University, Turku, Finland; Department of Cell and Developmental Biology, Feinberg School of Medicine, Northwestern University, Chicago, Illinois, USA; Max Perutz Labs, Medical University of Vienna, Vienna Biocenter Campus (VBC), Vienna, Austria; Turku Center for Disease Modeling, Turku, Finland; Turku Bioscience Centre, University of Turku and Åbo Akademi University, Turku, Finland

**Keywords:** Keratins, lamin, intermediate filament, colon epithelial cells, LINC proteins, proliferation, pRb, YAP

## Abstract

Keratin intermediate filaments (IFs) convey mechanical stability and protection against stress to epithelial cells, and may participate in nuclear structure and organization. Keratins are important for colon health as observed in keratin 8 knockout (K8^−/−^) mice, which exhibit colonic inflammation and epithelial hyperproliferation. Here, using a full body and two intestinal epithelial-specific K8^−/−^ knockout mouse models, we determine if cytoplasmic keratins affect the nuclear structure and lamina in epithelial colonocytes. K8^−/−^ colonocytes in vivo and in organoid cultures exhibit significantly decreased levels of the major lamins A/C, B1 and B2 in a colon-specific and cell-intrinsic manner independent of major changes in colonic inflammation or microbiota. Downregulation of K8 by siRNA in Caco-2 cells similarly decreases lamin A levels, which recover after re-expression of K8. K8 loss is associated with reduced plectin, LINC complex proteins and lamin-associated proteins, indicating a dysfunctional keratin-nuclear lamina coupling. Immunoprecipitation identifies complexes of colonocyte keratins with the LINC protein SUN2 and lamin A. Hyperphosphorylation of the lamin A-associated cell cycle regulator pRb in K8^−/−^ colonocytes together with increased nuclear localization of the mechanosensor YAP provide a molecular mechanism for the hyperproliferation phenotype. These findings identify a novel, colonocyte-specific role for K8 in nuclear function.

## Introduction

Keratins (K) are the most abundant intermediate filament (IF) proteins and divided into acidic type I keratins (K9-28) and basic/neutral type II keratins (K1-8 and K71-80) (Schweizer *et al*., 2006). Keratins form obligatory non-covalent heteropolymers in epithelial cells (Coulombe and Omary, 2002), and the main simple epithelial keratins in the colonic epithelium are K7 and K8 (type II), and K18, K19 and K20 (type I) (Zhou *et al*., 2003). Keratins have multiple functions, including providing cell stress protection, mechanical stability, protein and organelle scaffolding, and modulating protein targeting in cells and tissues (Toivola *et al*., 2005; Omary, 2017). Several diseases have been linked to keratin mutations, e.g. in skin and liver, while their role in colon and inflammatory bowel diseases (IBD) are still unclear (Coulombe *et al*., 1991; Ku *et al*., 2003; Toivola *et al*., 2015; Omary, 2017). However, K8 knockout (K8^−/−^) mice display a colonic phenotype marked by colitis, epithelial hyperproliferation, mistargeting and/or downregulation of membrane proteins (e.g. ion transporters) leading to diarrhea, deregulated epithelial differentiation and blunted colonocyte energy metabolism (Baribault *et al*., 1994; Toivola *et al*., 2004; Helenius *et al*., 2015; Asghar *et al*., 2016; Lähdeniemi *et al*., 2017). K8^+/−^ mice, which express 50 % less keratins than K8^+/+^ mice, exhibit an intermediate phenotype with moderate hyperproliferation, a partial ion transport defect, an increased susceptibility to experimental colitis, but no obvious spontaneous colitis under basal conditions (Asghar *et al*., 2015; Asghar *et al*., 2016; Liu *et al*., 2017). Interestingly, K8^−/−^ mice in two models of colorectal cancer (Misiorek *et al*., 2016), and K8^+/−^ in a model combining chemical colitis and colorectal cancer (Liu et al., 2017), are highly susceptible to induced colorectal cancer. The colitis and hyperproliferation in K8^−/−^ mice (Habtezion *et al*., 2005) and the tumor load in K8^+/−^ (Liu et al., 2017) can be partly ameliorated with antibiotics (Habtezion *et al*., 2005), indicating that the microbiota plays a crucial part in these K8-deficient colon phenotypes.

The nuclear lamina is a filamentous meshwork beneath the inner nuclear membrane (INM) that is composed of type V IF proteins, the nuclear A- and B-type lamins, and INM proteins (Wilson and Foisner, 2010). A-type lamins, mainly lamins A and C, are encoded from the LMNA gene through alternative splicing, and B-type lamins, primarily lamins B1 and B2, are encoded by the LMNB1 and LMNB2 genes, respectively (Dechat *et al*., 2008). Lamins are important for many nuclear processes, including maintenance of nuclear morphology and integrity, mechanotransduction, chromatin organization, gene expression, differentiation and proliferation (Dechat *et al*., 2010; Wilson and Foisner, 2010; Naetar *et al*., 2017; Brady *et al*., 2018; de Leeuw *et al*., 2018). These processes involve interactions between lamins and a multitude of lamin-binding proteins in the INM, nucleoplasm and chromatin, including LAP2-Emerin-MAN1 (LEM)-domain proteins, lamin B receptor (LBR) and SUN proteins (Wagner and Krohne, 2007; Wilson and Foisner, 2010). For example, a nucleoplasmic LEM-domain protein, lamina-associated polypeptide 2α (LAP2α), is crucial for maintaining and regulating a pool of nucleoplasmic lamin A/C (Naetar *et al*., 2008). LAP2α and nucleoplasmic lamin A/C interact with and promote the activity of the cell cycle repressor retinoblastoma protein (pRb), thus controlling and inhibiting cell proliferation (Naetar *et al*., 2008; Gesson *et al*., 2014; Vidak *et al*., 2018). Mutations in the lamin genes, mainly in the LMNA gene, cause many severe diseases termed laminopathies, such as Hutchinson-Gilford progeria syndrome (HGPS), muscular dystrophies and cardiomyopathies (Worman, 2012).

The nuclear lamina and the cytoskeleton are connected through LINC complexes, which are important for nuclear morphology, positioning and mechanotransduction (Crisp *et al*., 2006; Tapley and Starr, 2013; Osmanagic-Myers *et al*., 2015). LINC complexes are composed of outer nuclear membrane (ONM) KASH proteins, nesprins, that are connected to INM SUN proteins (Crisp *et al*., 2006). The cytoskeleton binds LINC complexes either directly or through cytolinker proteins, such as plectin, and, for example, epidermal keratin IFs in keratinocytes and mesenchymal vimentin IFs are connected to nesprin-3 through plectin (Wilhelmsen *et al*., 2005; Ketema *et al*., 2013; Almeida *et al*., 2015). Little is known about lamin and keratin synergy, however, epidermal K1/K10 loss has been linked to premature loss of nuclei, impaired nuclear integrity and decreased lamin A/C, emerin and SUN1 protein levels (Wallace *et al*., 2012), and K14 loss in epidermal keratinocytes leads to aberrations in the shape and size of the nucleus (Lee *et al*., 2012). Importantly, the role of simple-type epithelial keratins in regulating nuclear lamin function, especially in the colonic epithelium, is unknown (Brady *et al*., 2018). As we hypothesize that simple epithelial keratins may regulate lamins and the nucleus, the objective of this study was to assess if the loss of these keratins affect lamins and lamin-associated proteins in K8^−/−^ colonocytes. Here, we show for the first time that lamin protein levels correlate with keratin levels particularly in colonocytes, and that cytoplasmic keratins complex with lamins and couple to the nuclear lamina. Furthermore, the colon hyperproliferation observed in K8^−/−^ mice is a likely consequence of keratin-related changes in LINC complex and lamin-associated proteins.

## Results

### Loss of simple epithelial K8 correlates with decreased lamin levels in murine colonic epithelial cells *in vivo*

To determine whether simple epithelial keratins influence nuclear lamins in colonic epithelial cells, wee first analyzed lamin A, B and C levels in the colonic epithelium as a function of keratin levels using K8^−/−^ mice, in which most colonocyte keratins are depleted except for a thin keratin-layer at the apical membrane (Asghar *et al*., 2015).

Western blot analysis of crudely isolated colonic epithelial cells from K8^+/+^, K8^−/−^ and K8^+/−^ mice revealed significantly decreased protein levels of all major lamins, i.e. lamins A (2-fold), C (3-fold), B1 (5-fold) and B2 (10-fold), in K8^−/−^ mouse colonocytes compared to K8^+/+^ (Fig. 1A-F). K8^+/−^ colonocytes also exhibited significantly decreased lamin B2 protein levels (2-fold), but in contrast to K8^−/−^, the decrease was moderate (Fig. 1F). In addition, the levels of lamin A S22 phosphorylation (lamin A pS22) was decreased when adjusted for total lamin A protein levels in K8^−/−^ mouse colonocytes compared to K8^+/+^ (Fig. 1C). Lamin gene expression analysis by qRT-PCR showed that there was no significant change in lamin B1 mRNA levels, while lamin A/C and laminB2 mRNA levels were decreased on average by 50% in K8^−/−^ colonocytes compared to K8^+/+^ (Supplemental Fig. 1). Taken together, these data show that the protein levels of the major lamin isoforms A, C, B1 and B2 are decreased in K8^−/−^ colonocytes, to a lesser but generally intermediate extent in K8^+/−^ colonocytes, and that the downregulation is associated with decreased lamin gene expression, apart for lamin B1.

**Figure 1.**
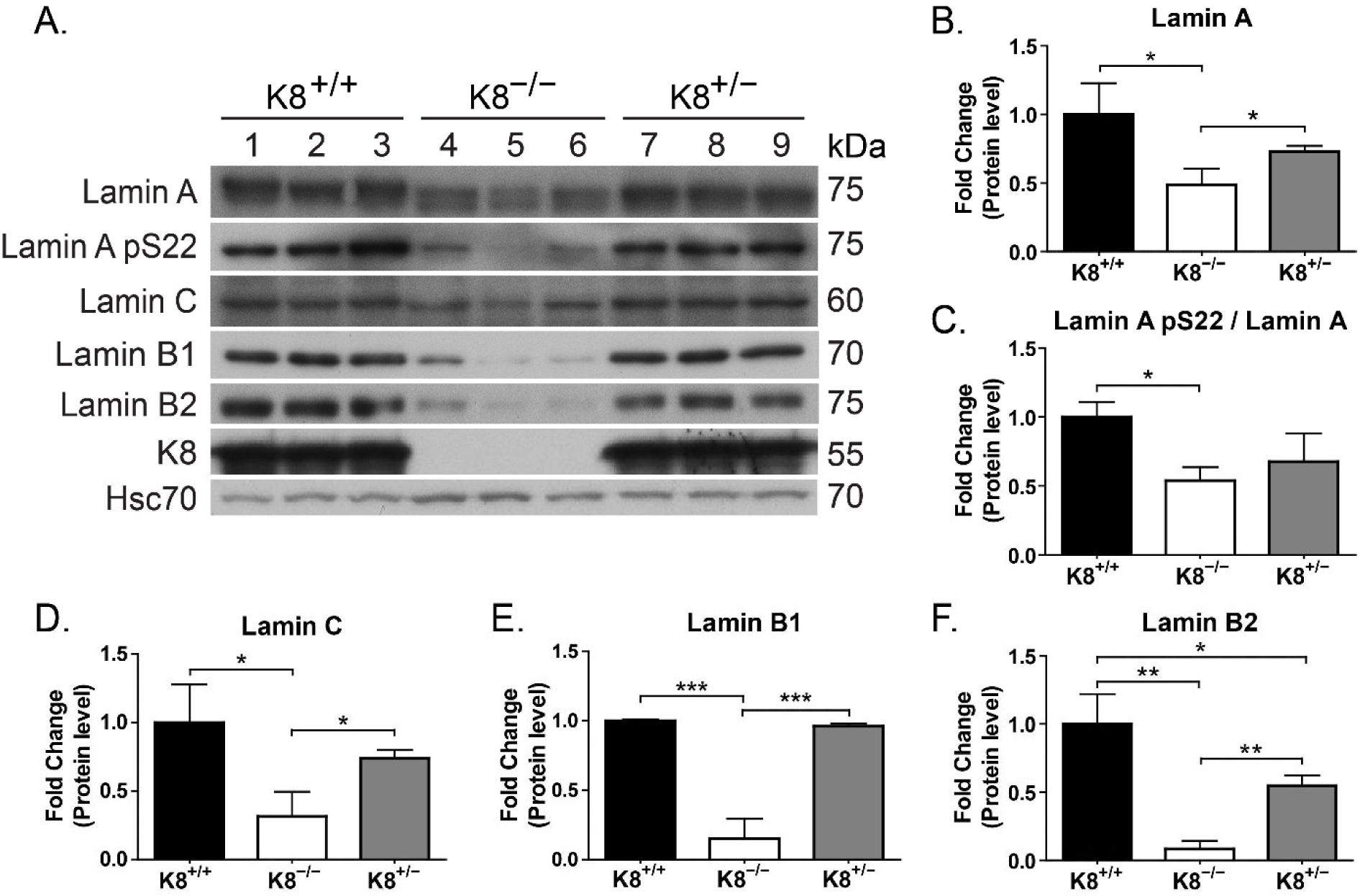
Lamins A, B1, B2 and C are downregulated in isolated K8^−/−^ mouse colon epithelial cells. A) Lysates of crudely isolated colon epithelium from K8^+/+^ (lanes 1-3), K8^−/−^ (lanes 4-6) and K8^+/–^ (lanes 7-9) mice (n=3) were immunoblotted for lamins A, lamin A pS22, lamin C, lamin B1, lamin B2, and K8. Hsc70 was used as a loading control. B-F) The immunoblots in A) were quantified and normalized to Hsc70 (or lamin A for lamin A pS22) protein levels. The results are representative of 2 other similar sets of mice and represent the mean (n=3) protein quantity ± SD with significant differences shown as * = p < 0.05, ** = p < 0.01 and *** = p < 0.001.

### Lamin downregulation in the absence of K8 is specific for epithelial cells in the colon

To carefully define the location of the decreased lamin levels seen in K8^−/−^ colon, lamin A immunostaining was performed on K8^+/+^ and K8^−/−^ colon. K8^+/+^ colonocytes express lamin A along the whole crypt, while clearly less lamin A was observed in the entire K8^−/−^ crypt compared to K8^+/+^ (Fig. 2A-B). Furthermore, K8^−/−^ lamin A levels were unaffected in non-epithelial cells of the lamina propria and the submucosa (Fig. 2A-B), showing that the loss of lamin occurs in the keratin-expressing epithelial cells.

**Figure 2.**
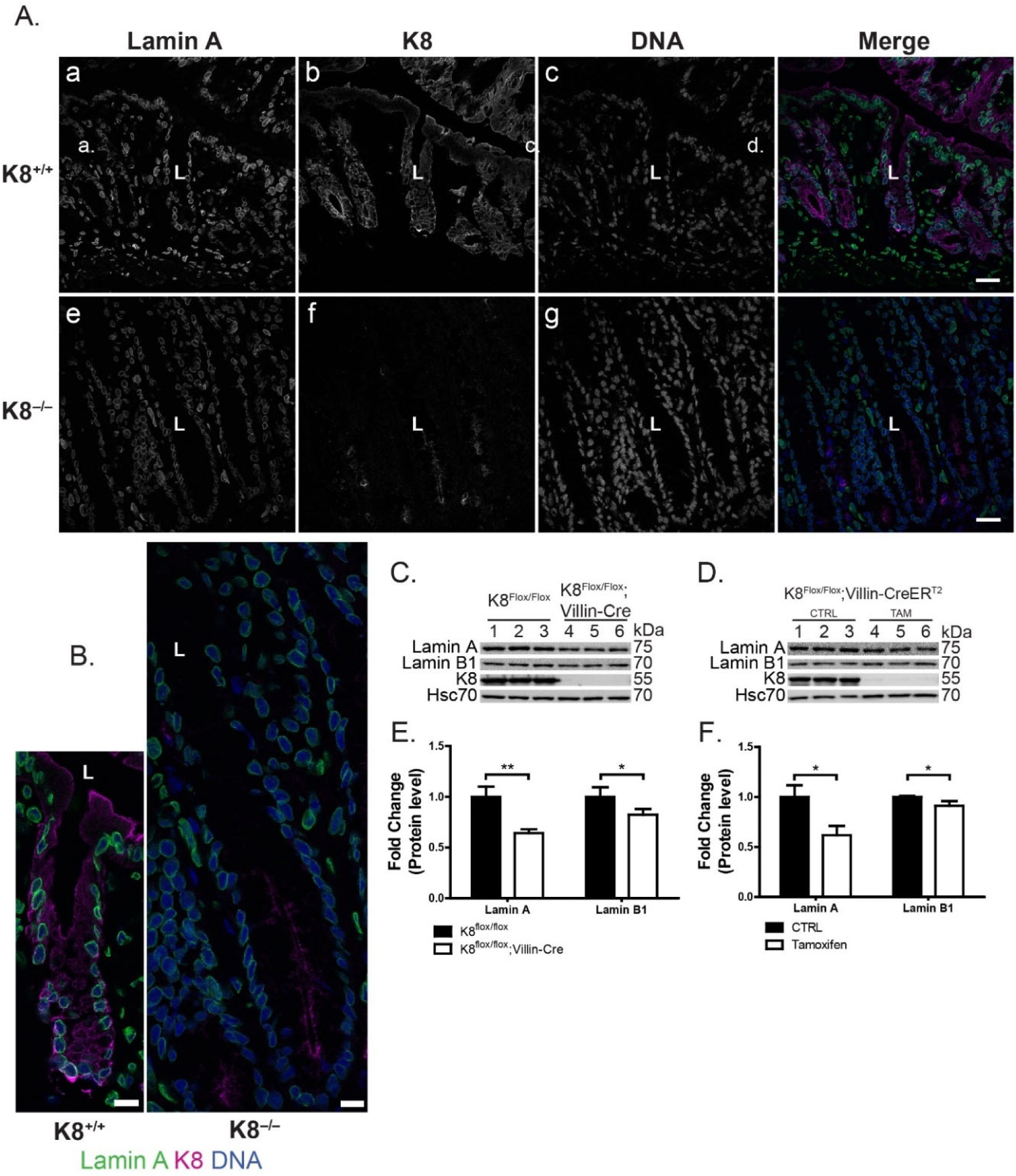
Loss of lamin A occurs only in colon crypt epithelial cells, and intestine-specific K8^−/−^ mouse colon epithelial cells have decreased lamin protein levels. A-B) K8^+/+^ and K8^−/−^ mouse colon tissue cryosections were immunostained for lamin A (green), K8 (magenta) and DNA (DRAQ5, blue). The individual images in A) and the enlarged merged pictures of crypts in B) show the colon epithelial cell-specific lamin A loss in the long and hyperproliferating crypts in the K8^−/−^ colon. Scale bars = 25 µm (A) and 10 µm (B). C-F) Lysates of crudely isolated colon epithelium from control K8^flox/flox^ (lanes 1-3 in C) and K8^flox/flox^;Villin-Cre (endogenous K8 knockdown; lanes 4-6 in C) mice and non-treated (control; CTRL; lanes 1-3 in D), and tamoxifen (TAM)-treated K8^flox/flox^;Villin-Cre-ER^T2^ (induced K8 knockdown; lanes 4-6 in D) mice were immunoblotted for lamins A and B1 and K8. The immunoblots were quantified and normalized to Hsc70 protein levels (loading control). The results represent the mean (n=3) protein quantity ± SD with significant differences shown as * = p < 0.05 and ** = p < 0.01.

We next asked if the loss of lamins in the keratin-deficient colon seen in the full K8^−/−^ mouse is colonocyte-autonomous, and for this colon-specific K8^−/−^ mice were established using the flox-Villin-Cre system. K8^flox/flox^ mice were bred with either Villin-Cre or Villin-CreEr^T2^ mice to generate K8 knockdown specifically in the intestinal epithelial cells (mucosal crypt layer). K8^flox/flox^;Villin-Cre mice are K8-deficient from birth, while the loss of K8 expression in K8^flox/flox^;Villin-CreER^T2^ mice is tamoxifen-inducible (1 injection/day for 5 days, whereafter mice were sacrificed after 25 days). In both colon-specific K8^−/−^ mouse models, lamin A and B1 protein levels were decreased in colonocytes *in vivo* (Fig. 2C-F and Supplemental Fig. 2A), implying that this phenotype is indeed colonocyte-intrinsic. Additionally, a similar reduction of lamin A/C protein levels, as seen *in vivo* (Fig. 1A, B and D), was observed *ex vivo* in K8^flox/flox^;Villin-Cre-ER^T2^ mouse colon 3D cultured organoids displaying an almost complete knockdown of K8 following tamoxifen-induced gene deletion (Supplemental Fig. 2B).

To assess if lamins are affected in other epithelial tissues normally expressing K8, lamin protein levels were analyzed in the small intestine (ileum), liver, pancreas and lung. In contrast to the K8^−/−^ colon, the small intestine in K8^−/−^ mice has a mild or negligible disease phenotype and does not display major hyperproliferation nor inflammation (Baribault *et al*., 1994; Ameen *et al*., 2001). The lamin A, B1/B2 and C protein levels in total lysates of the most distal part of the small intestine (ileum) were unchanged in K8^−/−^ mice compared to K8^+/+^ mice (Fig. 3). Similarly, no changes in lamin A levels were observed in total lysates of K8^−/−^ liver, pancreas and lung, while a decrease was observed in K8^−/−^ colon total lysates (Supplemental Fig. 3A-D). Taken together, these findings identify a colon and colonocyte-specific role for keratins in the regulation of lamin protein levels.

**Figure 3.**
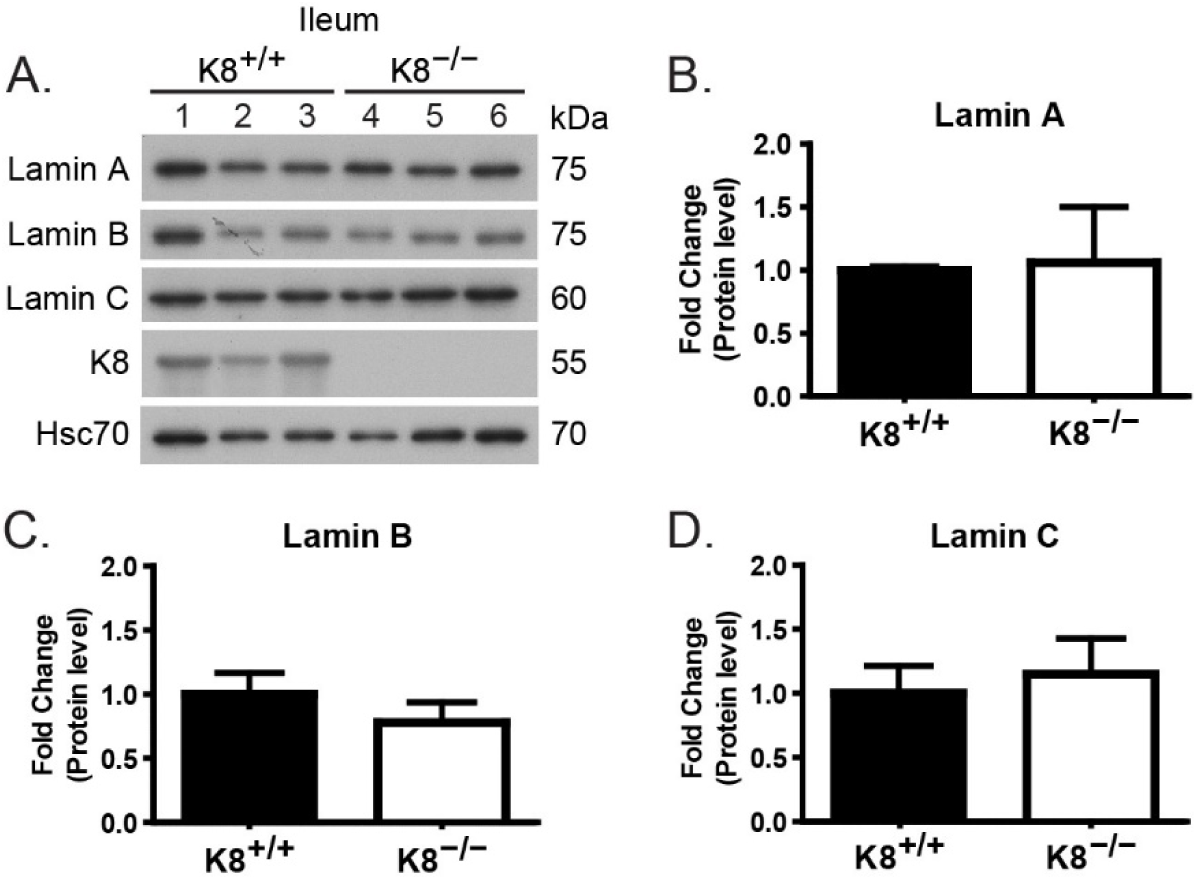
Lamin protein levels are comparable in K8^+/+^ and K8^−/−^ mouse ileum. A) Total lysates of ileum from K8^+/+^ (lanes 1-3) and K8^−/−^ (lanes 4-6) mice were immunoblotted for lamins A, B and C and K8. Hsc70 was used as a loading control. B-D) The immunoblots were quantified and normalized to Hsc70 protein levels. The results represent the mean (n=3) protein quantity ± SD. No statistically significant differences were observed between K8^+/+^ and K8^−/−^ samples.

### The colitis and microbiota levels do not contribute to lamin downregulation in K8^−/−^ mouse colon

The colonic microbiota plays a crucial role in the K8^−/−^ colon phenotype, as depletion of the colonic microbiota by broad-spectrum antibiotic treatment decreases inflammation and hyperproliferation in K8^−/−^ mice (Habtezion *et al*., 2005; Habtezion *et al*., 2011). Therefore, the impact of inflammation and microbiota on lamin protein expression was assessed in colonocytes from K8^+/+^ and K8^−/−^ mice treated with broad-spectrum antibiotics. Antibiotic treatment did not elicit any changes in, or normalization of, K8^−/−^ colonocyte lamin protein levels compared to control (baseline) mice (Fig. 4A). Decreased neutrophil infiltration in K8^− /−^ colon, as seen by myeloperoxidase staining, confirmed that the antibiotic treatment ameliorated the inflammation in K8^−/−^ colon (Fig. 4B) (Asghar *et al*., 2015). Moreover, lamin A, B1 and C protein levels remained unchanged in K8^+/+^ mice treated with the colitis-inducing chemical dextran sodium sulfate (DSS) compared to untreated mice (Supplemental Fig. 3E-H). These findings exclude the colonic microbiota and the colitis observed in K8^−/−^ colon as major contributing factors for the lamin changes and suggest a more direct role for keratins in the observed lamin phenotype.

**Figure 4.**
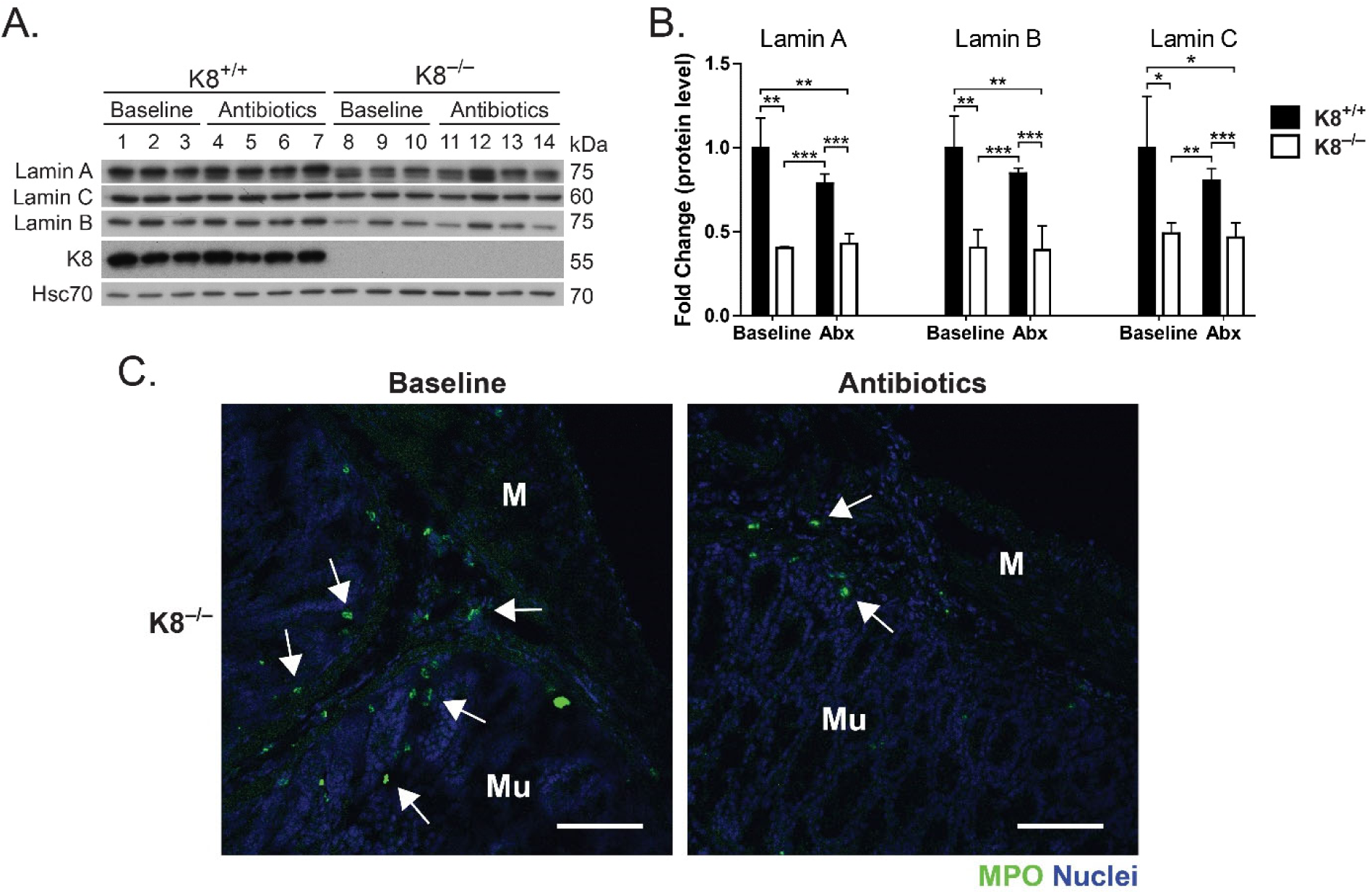
Absence of the colonic microbiota does not affect lamin protein levels in K8^+/+^ and K8^−/−^ mouse colon epithelial cells. A) Lysates of crudely isolated colon epithelium from non-treated (baseline) and antibiotic-treated (imipenem and vancomycin as described in Materials and methods) K8^+/+^ and K8^−/−^ mice were immunoblotted for lamins A, B and C and K8. Hsc70 was used as a loading control. B) The immunoblots were quantified and normalized to Hsc70 protein levels. The results represent the mean (n=3 for baseline and n=4 for antibiotic-treated mice) protein quantity ± SD with significant differences shown as * = p < 0.05, ** = p < 0.01 and *** = p < 0.001. C) The inflammatory status of the colon in baseline and antibiotic-treated K8^−/−^ mice was assessed by the presence of myeloperoxidase (MPO)-positive neutrophils (white arrows). Nuclei (blue) were visualized with DRAQ5. Scale bar = 100 μm; Mu, mucosa; M, muscularis.

### Keratins maintain lamin levels in colorectal adenocarcinoma cells

To assess whether the effect of K8 downregulation on colonocyte lamin levels *in vivo* is reproducible *in vitro*, human colorectal adenocarcinoma Caco-2 cells were subjected to sustained treatment with K8 and K18 siRNAs. Robust knockdown of keratins (about 80 %) elicited a significant decrease in lamin A (about 40 %) and lamin B1 (about 20 %) protein levels (Fig. 5A-E). Immunofluorescence staining of K8/K18 siRNA-treated cells showed a similar decrease of lamin A levels in cells lacking K8/K18, while the decrease in lamin B1 levels was not as obvious (Fig. 5F). This leads to an apparent shift in the lamin A/B ratio, with lamin B1 appearing more prominent than lamin A in cells lacking K8/K18, which is supported by the more robust keratin-dependent decrease of lamin A compared to lamin B1 in siRNA-treated cells (Fig. 5D-E). Importantly, allowing siRNA-treated cells to recover for 9 days after siRNA treatment completely normalized K8/K18 protein levels (Fig. 5 A-C), simultaneously rescuing the lamin A and B1 phenotype (Fig. 5B-E), indicating that lamin protein levels correlate with keratin levels in colon epithelial cells.

**Figure 5.**
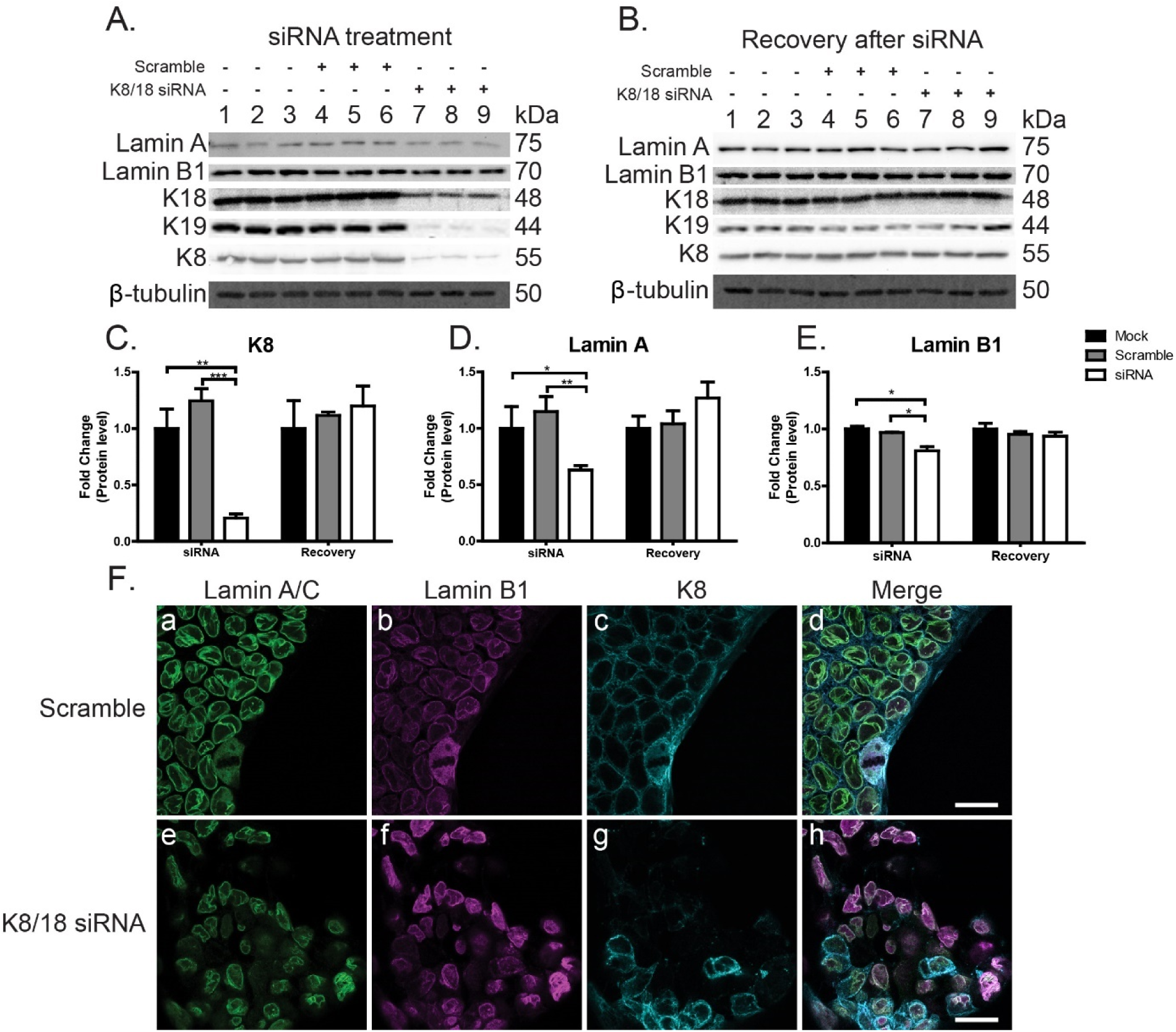
Caco-2 cells treated with K8/K18 siRNA exhibit reduced lamin protein levels, which can be rescued by re-expression of keratins. A) Mock-transfected (all reagents except siRNA), K8/K18 scramble siRNA-transfected and K8/K18 siRNA-transfected Caco-2 cells were immunoblotted for lamins A and B1, K8, K18 and K19. β-actin was used as a loading control. B) siRNA-treated cells were allowed to recover after siRNA-treatment and analyzed by immunoblotting for lamins A and B1, K8, K18 and K19. β-actin was used as a loading control. Recovery of the cells after siRNA-treatment led to re-expression of keratins and rescue of lamin expression. C-E) The immunoblots were quantified and normalized to β-actin and represent the mean (n=3) protein quantity ± SD with significant differences shown as * = p < 0.05, ** = p < 0.01 and *** = p< 0.001. F) K8/K18 scramble siRNA- and K8/K18 siRNA-treated Caco-2 cells were immunostained for lamin A (green), lamin B1 (magenta) and K8 (cyan). Scale bar = 25 µm.

### LINC complex protein levels are decreased in K8^−/−^ mouse colonocytes and correlate with increased nuclear translocation of YAP

Loss of K1/K10 disrupts nuclear integrity and may decouple the cytoskeleton and nucleus in the epidermis (Wallace *et al*., 2012). To assess whether nucleo-cytoskeletal coupling and nuclear integrity may be connected to K8 in colonocytes, the levels of the cytolinker protein plectin and several LINC complex and lamin-associated proteins were assayed. SUN1, SUN2 and the INM protein emerin had reduced gene expression in K8^−/−^ colonocytes as seen by qRT-PCR, while no change was seen in plectin or nesprin-3 mRNA levels (Supplemental Fig. 4A). However, the decrease in SUN1, SUN2 and emerin protein levels in K8^−/−^ colonocytes was larger than indicated by the qRT-PCR analysis, while also plectin protein levels were decreased significantly (Fig. 6A-E). Furthermore, SUN1 and emerin protein levels in K8^+/−^ colonocytes were moderately decreased compared to K8^+/+^ colonocytes (Fig. 6E). Additionally, SUN2 protein levels were unchanged in the small intestine (ileum) of K8^−/−^ mice compared to K8^+/+^ (Supplemental Fig. 4B), indicating that the keratin-associated effect on LINC and lamin-associated proteins is colon-specific.

**Figure 6.**
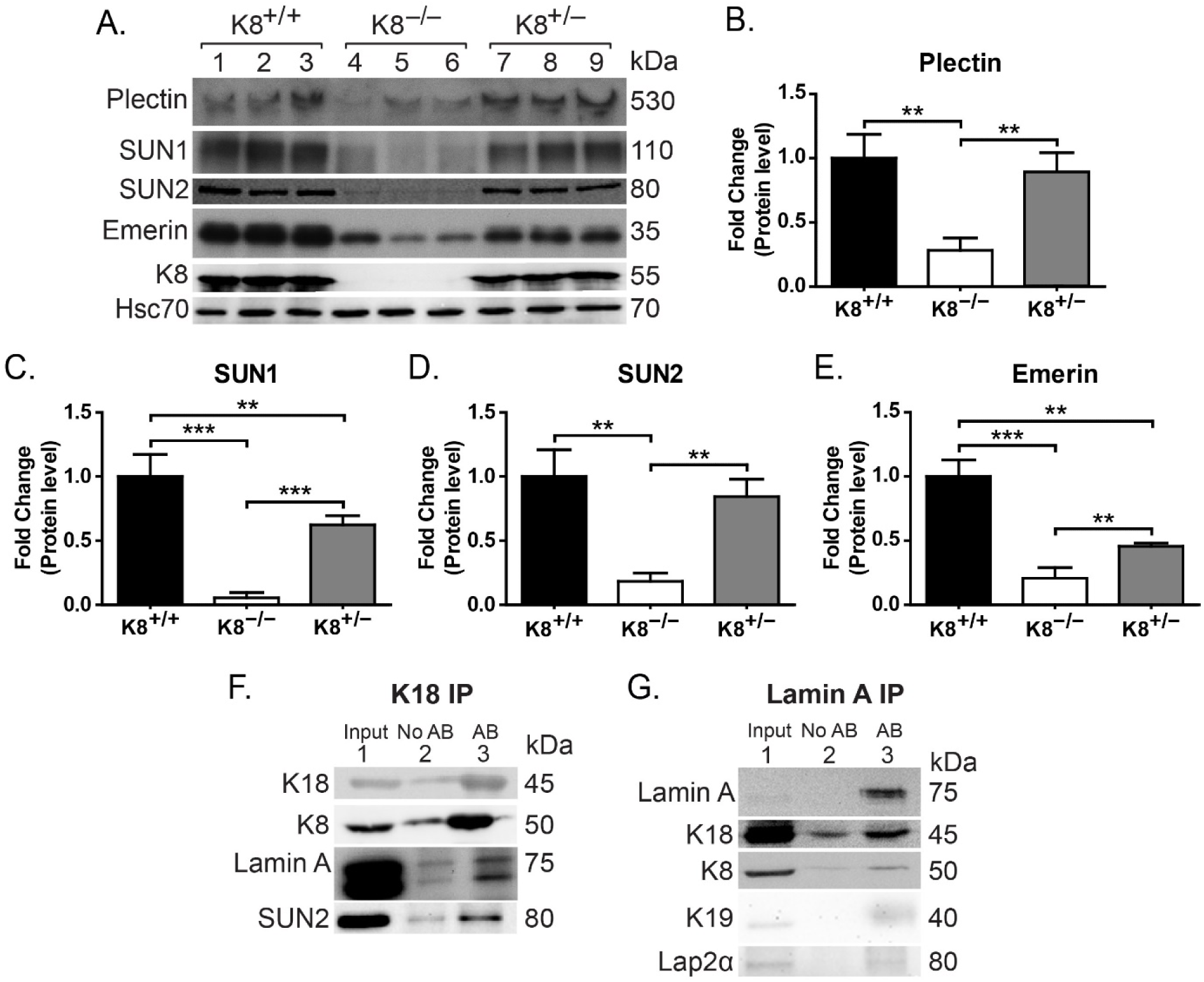
LINC protein levels are decreased in K8^−/−^ colon epithelial cells. A) Lysates of crudely isolated colon epithelium from K8^+/+^ (lanes 1-3), K8^−/−^ (lanes 4-6) and K8^+/−^ (lanes 7- 9) mice (n=3) were immunoblotted for plectin, nesprin-3, SUN1, SUN2, emerin and K8. Hsc70 was used as a loading control. B-E) The immunoblots were quantified and normalized to Hsc70 protein levels and represent the mean (n=3) protein quantity ± SD with significant differences shown as * = p < 0.05, ** = p < 0.01 and *** = p < 0.001. F-G) Caco-2 cell lysates (input) were used in immunoprecipitation assays where K8/K18 (F) or lamin A (G) were immunoprecipitated using K18 or lamin A antibodies, respectively. Immunoprecipitates were immunoblotted for K8, K18, K19, lamin A, SUN2 and LAP2α. K8/K18 co-immunoprecipitated lamin A and SUN2, and lamin A co-immunoprecipitated K8, K18, K19 and LAP2α. No AB indicates a negative control sample, in which the immunoprecipitation step was performed without antibody.

To understand the molecular connection between cytoplasmic keratins and nuclear lamins, K8/K18 or lamin A co-immunoprecipitation assays were performed on lysates of Caco-2 cells. K8/K18 co-immunoprecipitated with lamin A and the LINC protein SUN2 while, conversely, lamin A co-immunoprecipitated K8, K18 and K19 (Fig. 6F-G). These data indicate that keratins can complex with LINC complexes and lamins, suggesting that keratins could physically link to the lamina via the LINC complex and may thus affect the mechanical properties of the nucleus and nuclear mechanotransduction, and thereby modulate nuclear function.

The mechanosensitive transcriptional regulator Yes-associated protein (YAP) has been implicated as an important integrator of mechanotransduction, proliferation as well as regeneration in the colon, and the loss of YAP impairs colon tissue regeneration upon injury (Cai *et al*., 2010; Yui *et al*., 2018). As mechanical regulation of YAP involves the actomyosin cytoskeleton, Src family kinases and intact LINC complexes (Elosegui-Artola *et al*., 2017; Ege *et al*., 2018), and LINC complex and LINC complex-associated proteins important for mechanotransduction are downregulated in K8^−/−^ colonocytes (figure 6) (Lombardi and Lammerding, 2011), the cellular localization and nuclear translocation of YAP was investigated to test whether the keratin cytoskeleton may be involved in mechanical regulation. In K8^+/+^colonocytes, YAP levels were higher in the crypt bottom (Fig. 7A), where lamin levels are relatively low (Fig. 2A-B), compared to higher up in the crypt. Interestingly, K8^−/−^ colonocytes, which have decreased lamin levels (Fig. 1) and are nearly keratin-free (Asghar *et al*., 2015), showed a clearly increased nuclear translocation of YAP compared to K8^+/+^ colonocytes (Fig. 7A), while YAP protein levels were comparable in K8^+/+^ and K8^−/−^ colonocytes. *In vitro*, Caco-2 cells treated with K8/K18 siRNA exhibit decreased cytoplasmic YAP compared to mock-transfected (data not shown) or K8/K18 scramble siRNA-transfected cells, while the levels of nuclear YAP appears similar or slightly elevated in K8/K18 siRNA-treated cells compared to controls (Fig. 7B), supporting the *in vivo* data. Interestingly, similar to K8/K18 scramble siRNA-transfected cells, the few K8-retaining cells in K8/K18 siRNA-treated cell cultures show higher cytoplasmic YAP compared to cells lacking K8 (Fig. 7B). These results support the notion that keratins may be involved in the mechanical coupling between the cytoskeleton and the nucleus in colonic epithelial cells.

**Figure 7.**
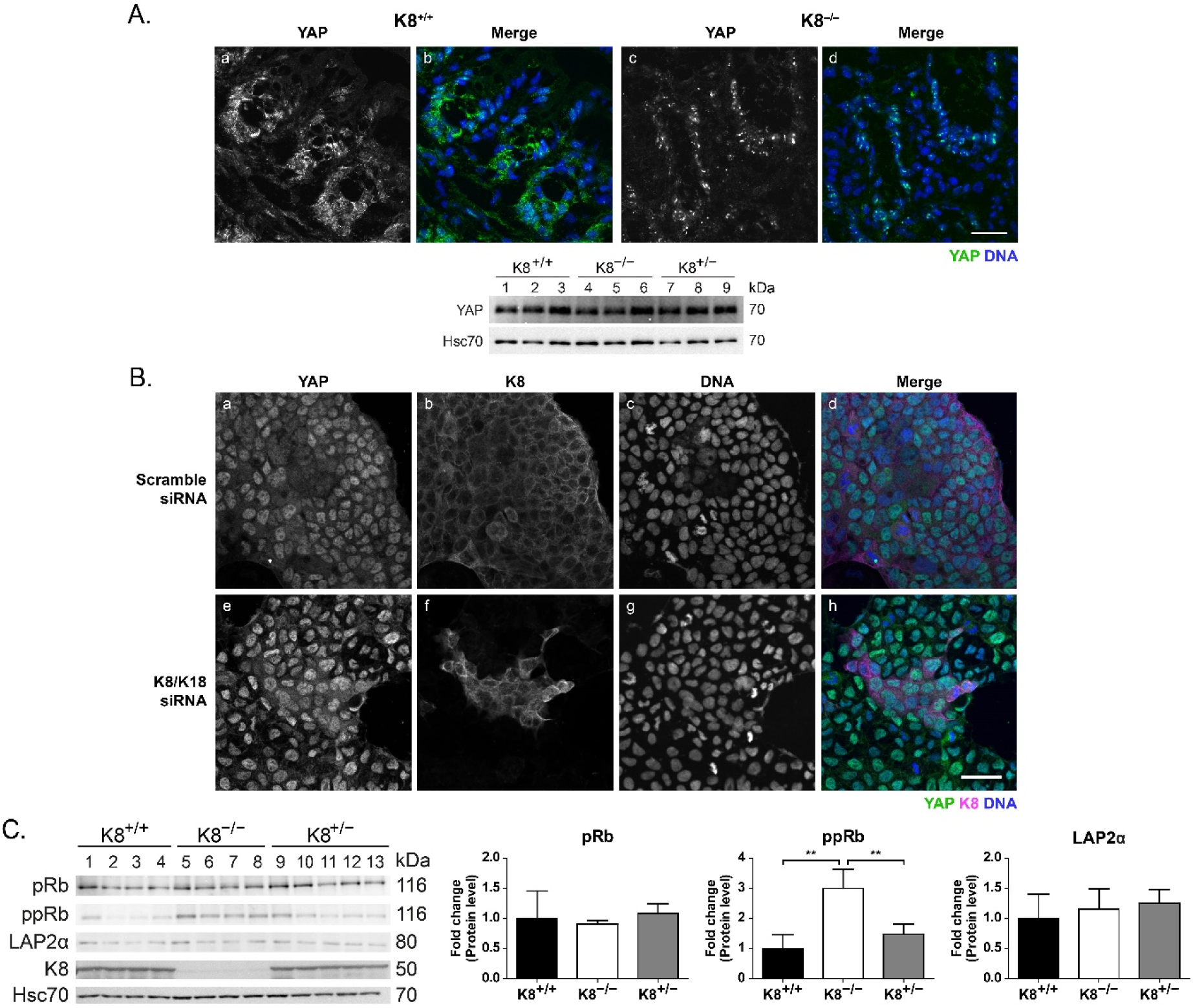
Loss of K8 correlates with increased nuclear translocation of YAP and hyperphosphorylation of pRb in colon epithelial cells. A) K8^+/+^ and K8^−/−^ mouse colon tissue cryosections were immunostained for YAP (green) and DNA (DRAQ5, blue). Scale bar = 25 µm. Lysates of crudely isolated colon epithelium from K8^+/+^ (lanes 1-3), K8^−/−^ (lanes 4-6) and K8^+/–^ (lanes 7-9) mice (n=3) were immunoblotted for YAP. Hsc70 was used as a loading control. B) Caco-2 cells treated with K8/K18 scramble siRNA (negative control) or K8/K18 siRNA were immunostained for YAP (green), K8 (magenta) and DNA (DRAQ5, blue). Scale bar = 50 µm. C) Lysates of crudely isolated colon epithelium from K8^+/+^ (lanes 1-4), K8^−/−^ (lanes 5-8) and K8^+/−^ (lanes 9-13) mice were immunoblotted for pRb, phosphorylated pRb (ppRb; phosphor-Ser 807 and phosphor-Ser 811), LAP2α, and K8. Hsc70 was used as a loading control. The immunoblots were quantified and normalized to Hsc70 protein (or against pRb for ppRb) levels and represent the mean protein quantity ± SD with significant differences shown as ** = p < 0.01.

### Lamin-associated protein pRb is hyperphosphorylated in K8^−/−^ mouse colonocytes, implying a disturbance in cell cycle regulation

We next addressed the mechanism involved in linking K8 and proliferation. Since changes in lamin B1 levels have previously been associated with the regulation of proliferation and senescence (Shimi *et al*., 2011; Barascu *et al*., 2012; Freund *et al*., 2012), the prevalence of senescence in the colon was analyzed using senescence-associated β-galactosidase staining in both proximal colon (PC) and distal colon (DC) of K8^+/+^ and K8^−/−^ mice. In both K8^−/−^ PC and DC the number of senescent cells was decreased compared to K8^+/+^ mice (Supplemental Fig. 5), indicating that the keratin-loss dependent lamin B1 reduction is clearly not increasing senescence.

A-type lamins bind LAP2α, and together they interact with and promote pRb-mediated inhibition of cell cycle progression, and consequently, proliferation (Naetar et al., 2008). As K8^−/−^ mouse colon is characterized by hyperproliferation (Baribault et al., 1994), and lamins A/C are downregulated in K8^−/−^ colonocytes (Fig. 1), K8^+/+^, K8^−/−^ and K8^+/−^ colonocytes were analyzed for the protein levels of LAP2α and pRb, and the levels of phosphorylated pRb (phosphor-Ser 807 and phoshpo-Ser 811), which constitutes the inactive form of pRb (Weinberg, 1995; Gesson *et al*., 2014). No significant changes in LAP2α nor pRb protein (Fig. 7C) or LAP2α mRNA levels (Supplemental Fig. 4A) were seen in K8^−/−^ colonocytes, however, pRb was highly hyperphosphorylated (3-fold compared to K8^+/+^ and K8^+/−^ colonocytes when normalized to pRb) (Fig. 7C), indicating that pRb may be inactivated in K8^−/−^ colonocytes. Since keratins affect lamin and lamin-associated protein levels, we tested if the reverse relationship occurs. Analysis of LAP2α^−/−^ mice, which exhibit epithelial hyperproliferation and increased pRb phosphorylation in the skin, and an increased amount of proliferating transit amplifying cells in the colon (Naetar *et al*., 2008), showed no change in either keratin nor lamin protein levels in LAP2α^−/−^ colonocytes (Supplemental Fig. 4C). This suggests that the complex containing lamin A/C, LAP2α and pRb is downstream of keratin-lamin interaction, and hence, loss of LAP2α does not alter keratin protein levels.

## Discussion

In this study, we demonstrate for the first time to our knowledge a molecular link between cytoplasmic simple epithelial keratins (K8 and its main partners K18 and K19), LINC complex proteins, lamin-associated proteins and nuclear lamins in colonocytes (summarized in Fig. 8). We show that the absence of keratins in colon epithelial cells leads to significantly decreased protein levels of the major lamin isoforms A, C, B1 and B2 *in vivo* in K8^−/−^ mice. Similarly, lamin A and B1 protein levels decrease *in vitro* in human Caco-2 cells treated with K8/K18 siRNA, and *ex vivo* in K8^flox/flox^ Villin-Cre-ER^T2^ 3D colon organoids. Reduced lamin gene expression, except for lamin B1, is likely the cause for the decreased lamin protein levels observed in K8^−/−^ colon, however, it is unclear how the loss of keratins could affect lamin expression, as little is known about lamin gene regulation (Dechat *et al*., 2008). Interestingly, the decrease in lamin protein levels was colon-specific, as several other K8^−/−^ simple epithelial tissues, where K8 is a major or the only type II keratin, exhibited normal lamin levels, including normal SUN2 protein levels as assessed for the small intestine. The correlation between keratin loss and decreased lamin levels is likely independent of the microbiota or colitis observed in K8^−/−^ mice, as antibiotic treatment, which eliminates the colonic microbiota and ameliorates the colitis and hyperproliferation (Habtezion *et al*., 2011), did not normalize lamin levels. Very little is known regarding the potential roles of lamins in colon homeostasis and function (Brady *et al*., 2018). Low levels of A-type lamins correlate with increased gastric polyp size (Wang *et al*., 2015), and loss of A-type lamins in stage II and III colorectal cancers (CRC) puts patients at severe risk of cancer resurgence (Belt *et al*., 2011). Since the lack or decreased levels of colonic keratins leads to hyperproliferation and increased susceptibility to CRC tumorigenesis (Misiorek *et al*., 2016; Liu *et al*., 2017), the lamin phenotype in these mice may be a contributing factor toward the pro-tumorigenic phenotype. Nevertheless, intestinal epithelial-specific lamin A knockdown in mice did not lead to increased proliferation as measured by ki67 (Wang *et al*., 2015).

**Figure 8.**
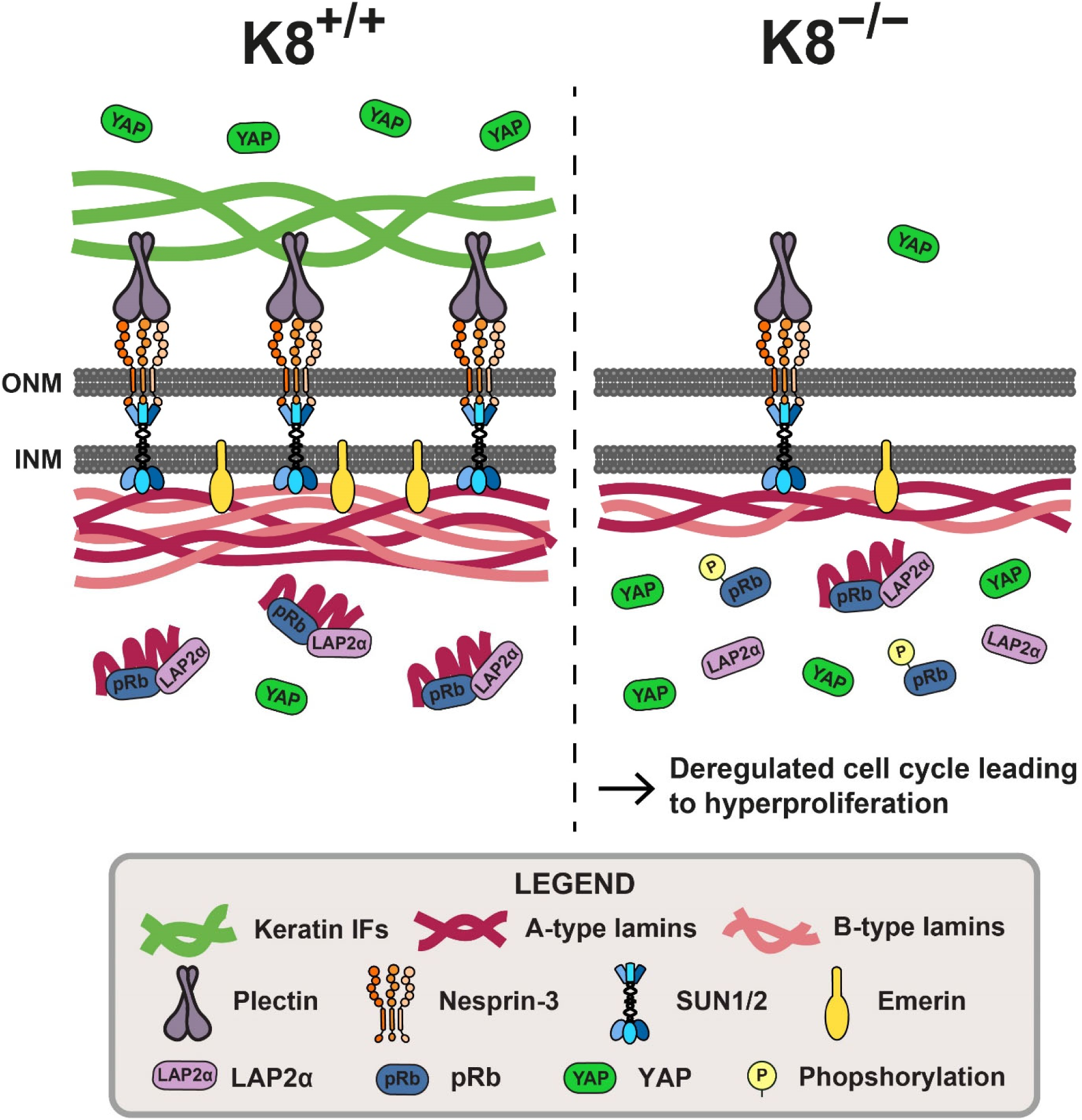
Cytoplasmic simple epithelial keratins complex with nuclear lamins, LINC complex proteins and lamin-associated proteins, maintaining keratin-nuclear lamina coupling and proliferation homeostasis in colonic epithelial cells. In this summary schematic, the findings of this study are summarized. In healthy colon epithelial cells (K8^+/+^), cytoplasmic K8-containing keratin IFs couple with and/or stabilize plectin, LINC complex proteins, lamins and lamin-associated proteins, thus helping maintain nucleocytoskeletal coupling and nuclear function. Loss of cytoplasmic keratins (K8^−/−^) in colon epithelial cells correlates with decreased plectin, LINC complex protein, lamin and lamin-associated protein levels, which likely disrupts these nuclear membrane complexes. Significantly, the decreased A-type lamin levels in K8^−/−^ colonocytes correlates with, and allows for, increased hyperphosphorylation of pRb, which in turn promotes cell cycle progression. Furthermore, K8-deficient colonocyte nuclei exhibit increased accumulation of the mechanosensor YAP. Together, these changes may contribute to the hyperproliferation phenotype observed in K8^−/−^ mouse colon and define a role for cytoplasmic keratins in nuclear integrity and function.

Lamin-associated proteins depend on A-type lamins, and vice versa, for stabilization and correct positioning in the nucleus (Vaughan *et al*., 2001; Libotte *et al*., 2005; Naetar *et al*., 2008; Muchir *et al*., 2009). Our data show that K8^−/−^ mouse colonocytes have decreased levels of emerin, which is anchored to the inner nuclear membrane by lamin A, and therefore the absence of emerin could be caused by decreased lamin A protein levels (Vaughan *et al*., 2001; Muchir *et al*., 2009). LAP2α is essential for maintaining a nucleoplasmic pool of A-type lamins (Dechat *et al*., 2000; Naetar *et al*., 2008), and together with lamin A/C stabilizes and promotes the activity of the important cell cycle regulator pRb (Markiewicz *et al*., 2002; Naetar *et al*., 2008; Gesson *et al*., 2014). In the present study, LAP2α and pRb levels were comparable in K8^+/+^ and K8^−/−^ colonocytes, while, in contrast, pRb was hyperphosphorylated in K8^−/−^ colonocytes. pRb-mediated cell cycle regulation is phosphorylation-dependent, as active, hypophosphorylated pRb inhibits cell cycle progression, while hyperphosphorylation leads to its inactivation, causing increased cell cycle progression and proliferation (Weinberg, 1995; Gesson *et al*., 2014). Thus, the increased hyperphosphorylation of pRb in K8^−/−^ colonocytes could be a major contributing factor to the hyperproliferation observed in K8^−/−^ mice (Baribault *et al*., 1994; Naetar *et al*., 2008; Gesson *et al*., 2014). Intriguingly, similar to K8^−/−^ mice (Lähdeniemi *et al*., 2017), LAP2α^−/−^ mice were reported to exhibit an upregulation of proliferating transit amplifying cells and enhanced colonic proliferation caused by LAP2α-loss-mediated depletion of nucleoplasmic A-type lamins and inactivation of pRb through phosphorylation (Naetar *et al*., 2008). However, we observed no major changes in keratin or lamin levels in LAP2α^−/−^ colonocytes (Supplemental Fig. 4C). Similarly, the loss of A-type lamins in mice causes mild epidermal hyperplasia comparable to that seen in LAP2α^−/−^ mice (Naetar *et al*., 2008), and pRb loss in mice leads to hyperproliferation and stimulates the differentiation of enterocytes, goblet cells, enteroendocrine cells and Paneth cells in the small intestine (Yang and Hinds, 2007). While we cannot rule out a more direct role of K8 in nuclear function similar to K17, which was recently shown to enter the nucleus where it can modulate several signaling pathways (Escobar-Hoyos *et al*., 2015; Hobbs *et al*., 2015), our findings imply that keratins act upstream of, and help stabilize, lamins and lamin-associated proteins in the nuclear lamina. Furthermore, these findings offer a first potential mechanism for the hyperproliferation and increased susceptibility to colorectal cancer caused by K8 deletion in the colonic epithelium.

Weakening or loss of the connection between the nuclear lamina and the cytoplasmic cytoskeleton can be caused by the loss and/or mislocalization of lamins and/or LINC complex proteins, leading to the loss of anchoring sites for connecting the nuclear lamina to LINC complexes (Lombardi and Lammerding, 2011). As several LINC complex proteins, including SUN1 and SUN2, and the major lamin isoforms are downregulated in K8^−/−^ colonocytes, we can surmise that coupling of LINC complexes and the nuclear lamina may be compromised in these cells. While we were unable to probe nesprin-3 protein levels due to the lack of a good antibody, nesprin-3 gene expression was unaffected in K8^−/−^ colonocytes. Nonetheless, as SUN1 and SUN2 are downregulated in K8^−/−^ colonocytes, and SUN proteins are required for correct localization of nesprin-3 at the ONM (Ketema *et al*., 2007), nesprin-3 is presumably displaced from the ONM. Furthermore, plectin, a cytolinker protein known to link nesprin-3 to vimentin and keratins (Wilhelmsen *et al*., 2005; Ketema *et al*., 2013; Almeida *et al*., 2015), is decreased in K8^−/−^ colonocytes, which supports the notion that the nuceloskeleton-cytoskeleton link is disrupted from the outside of the nucleus due to the loss of keratins and plectin. Our data in colonocytes support the findings reported for other cell types, that a continuous coupling between the cytoplasmic cytoskeleton and the nuclear lamina and the presence of lamin-associated proteins, such as emerin, are important for maintaining nuclear integrity and mechanosignaling (Lombardi and Lammerding, 2011; Osmanagic-Myers *et al*., 2015). Due to the decrease or loss of many proteins along this axis, it is highly likely that nuclear integrity and mechanosignaling is weakened or disrupted in K8^−/−^ colonocytes, and that nuclei in these cells are more vulnerable to mechanical stress. This is supported by the findings that epidermal K1/K10 loss in mice is associated with decreased levels of lamin A/C, emerin and SUN1 and a likely decoupling of the cytoskeleton and the nuclear lamina, leading to weakened nuclear integrity and premature loss of nuclei (Wallace *et al*., 2012). Similarly, mesenchymal vimentin IFs were recently shown to mechanically support nuclei, as the loss of vimentin caused increased nuclear rupture and DNA damage during cell migration (Patteson *et al*., 2019). Interestingly, in the present study, we observed increased nuclear translocation of the mechanosensor and transcriptional regulator YAP in K8^−/−^ compared to K8^+/+^ colonocytes, implying YAP activation by mechanical cues (Piccolo *et al*., 2014; Yui *et al*., 2018). In addition, K8/K18 siRNA-treated Caco-2 cells exhibit a robust nuclear YAP staining and decreased cytoplasmic YAP, compared to keratin-expressing cells. YAP activity was reported nonessential for intestinal homeostasis (Cai *et al*., 2010; Azzolin *et al*., 2014), however, upon DSS-induced colitis, YAP activity induced through mechanotransduction is indispensable for proliferation and regeneration in damaged colonic epithelium (Cai *et al*., 2010; Yui *et al*., 2018). YAP activation by mechanical cues involves the actomyosin cytoskeleton, Src family kinases and mechanical coupling of the cytoskeleton and nucleus (Elosegui-Artola *et al*., 2017; Ege *et al*., 2018), which would suggest that accumulation of nuclear YAP should be perturbed in K8^− /−^ colonocytes, as these cells exhibit reduced LINC complex proteins (Fig. 6) and absent lateral and patchy apical F-actin (Toivola *et al*., 2004). As nuclear translocation of YAP is possible in K8^−/−^ colonocytes, the few remaining LINC proteins may form functional units which facilitate YAP translocation. Alternatively, cytoplasmic keratin filaments may be necessary for limiting nuclear translocation of YAP, which needs to be further investigated.

Mechanically-induced nuclear and transcriptionally active YAP promotes cell proliferation in many cell types, including epithelial cells (Piccolo *et al*., 2014). As YAP is important for repairing epithelium in the colon upon DSS-induced colitis (Cai *et al*., 2010; Yui *et al*., 2018), and K8^−/−^ mice develop colitis, it could be surmised that a YAP-induced regenerative state in K8^−/−^ colon epithelium contributes to the hyperproliferation observed in K8^−/−^ colon. The heightened regenerative and proliferative state in these mice is also evident from increased activation of IL-22 and STAT3 signaling (Misiorek *et al*., 2016). Little is known about the role of IFs or keratins in the regulation of YAP. However, the “rim-and-spoke” hypothesis for IFs (Quinlan *et al*., 2017), in which radial keratin spokes connecting the plasma membrane with the ONM could potentiate a mechanosensory function for keratins, thus fulfilling a similar or complementary function in mechanosensing for IFs similar to actin filaments and stress fibers.

Lamin B1 loss or upregulation have been associated with cellular senescence, either as causes or markers of senescence both *in vitro* (e.g. human fibroblasts or keratinocytes) (Shimi *et al*., 2011; Barascu *et al*., 2012; Freund *et al*., 2012; Dreesen *et al*., 2013) and *in vivo* (e.g. mouse liver) (Freund *et al*., 2012; Dreesen *et al*., 2013). While decreased lamin B1 levels have been linked to senescence, it has also been linked to increased apoptosis (Harborth *et al*., 2001). Surprisingly, in disagreement with these findings, K8^−/−^ colonocytes exhibiting decreased lamin B1 levels show decreased senescence and apoptosis, and increased proliferation (Baribault *et al*., 1994; Habtezion *et al*., 2011; Lähdeniemi *et al*., 2017). Interestingly, the decreased senescence could be explained by the increased hyperphosphorylation of pRb in K8^−/−^ colonocytes, as senescent cells are characterized by a lack of pRb phosphorylation (Stein *et al*., 1990; Gire and Dulic, 2015) and senescence can be bypassed by pRb inactivation (Sage *et al*., 2003).

In conclusion, we show in this study that simple epithelial keratin filaments offer support to lamins, LINC complex proteins and lamin-associated proteins in colonic epithelial cells, and that K8 loss leads to dramatically decreased levels of these proteins. Consequently, the coupling between the cytoskeleton and nuclear lamina may be disrupted. Inside the nucleus, increased pRb hyperphosphorylation likely deregulates cell cycle regulation and inhibits senescence, while nuclear translocation of YAP is increased, and collectively these changes likely contribute to the hyperproliferation in K8^−/−^ colon. Our findings indicate a novel, colonocyte-specific role for K8 in maintaining lamin levels and nuclear function, and that K8 may be a novel cytoskeletal factor involved in the mechanical coupling of the cytoskeleton and the nucleus.

## Materials and methods

### Experimental animals and sample collection

K8^+/+^, K8^+/−^ and K8^−/−^ mice in the FVB/n background (Baribault *et al*., 1994) were bred and genotyped by PCR as previously described (Baribault *et al*., 1994). All animals were housed at the Central Animal Laboratory of the University of Turku and treated according to animal study protocols (nos. 197/04.10.07/2013 and 3956/04.10.07/2016) approved by the State Provincial Office of South Finland. The mice were euthanized by CO_2_ inhalation and tissue samples were collected. Pieces of proximal colon (PC) and distal colon (DC) were collected for protein and RNA analysis, immunofluorescence staining and senescence detection. Colon epithelium was collected for protein and RNA analysis by scraping the luminal side of the colon with a chilled glass slide as previously described (Helenius *et al*., 2015). Ileum, liver, pancreas and lungs were collected for protein analysis. Protein analysis of colon epithelium and PC were performed on samples from 2–5 months old male and female mice unless otherwise stated. Pancreas were collected from 2–3 months old male and female mice, lungs were collected from 8–10.5 months old male and female mice, and liver and ileum were collected from 5.5–6.5 months old male and female mice. K8^flox/flox^ Villin-Cre mice in the C57BL/6 background were of mixed gender and 4 months of age. LAP2α mice were of the C57BL/6 background and 3–3.5 months of age.

### Tamoxifen treatment of mice

A 15 mg/ml solution of tamoxifen (Sigma Aldrich, CA, USA) was prepared by dissolving 30 mg tamoxifen in 0.2 ml pure EtOH and then diluting it with 1.8 ml corn oil (Sigma Aldrich, CA, USA). Vehicle substance was prepared by mixing 0.2 ml EtOH with 1.8 ml corn oil. K8^flox/flox^ Villin-Cre-ER^T2^ mice in the C57BL/6 background received an intraperitoneal injection with 100 µl of either tamoxifen (1.5 mg tamoxifen/mouse) or vehicle solution once per day for five consecutive days, whereafter the mice were kept for 3.5 weeks before they were sacrificed.

### Antibiotic treatment of mice

Male and female K8^+/+^ and K8^−/−^ mice were treated with the broad-spectrum antibiotics vancomycin and imipenem (Hospira, IL, USA) administered via drinking water (68 mg/kg body weight/day of each antibiotic) for 8 weeks starting at 18-19 days of age, while control mice received normal drinking water. The drinking water (with or without antibiotics) was changed three times a week. Upon completion of the antibiotic treatment, the 2.5 months old mice were euthanized by CO_2_ inhalation, and colon epithelium was collected as described above.

### DSS treatment of mice

2 % DSS (40 kDa, TdB Consultancy AB, Sweden) was administered in autoclaved water to 2.5 months old male BALB/c mice for 8 days. Upon completion of the DSS treatment, the mice were euthanized by CO_2_ inhalation and colon total lysate samples were collected.

### Organoid culture

Colon organoids from K8^flox/flox^;Villin-Cre-ER^T2^ mice were isolated according to a modified previously described method (Stemcell Technologies, UK). Briefly, the mice were euthanized by CO_2_ inhalation and the colon was collected, cut open, washed with ice-cold PBS and incubated in 30 ml of 5 mM ethylenediaminetetraacetic acid (EDTA; pH 8) in PBS under rotation at RT for 20 minutes. Next, the colon was rinsed with PBS, followed by vigorous shaking by hand to mechanically detach the crypts. Isolated crypts were plated at approximately 2 crypts/μl in Matrigel (Corning, NY, USA) domes diluted 1:1 with advanced DMEM/F12 media (Stemcell Technologies, UK), with 50 μl domes per well. Once polymerized, 500 μl of IntestiCult Organoid Growth Medium (Stemcell Technologies, UK) supplemented with 10 µM Y-27632 (Adooq Bioscience, CA, USA) was added to each well and the colon organoids were cultured at 37 °C in a 5 % CO_2_ atmosphere for 48 hours. After the initial 48-hour incubation with medium supplemented with Y-27632, the organoids were incubated with medium without Y-27632 and the medium was exchanged every 48 hours. Colon organoids were harvested for immunoblotting by dissolving the Matrigel with 500 µl Gentle Cell Dissociation Reagent (Stemcell Technologies, UK) for 1 minute, whereafter the samples were centrifuged for 5 minutes at 2000 g. The pellet was homogenized with homogenization buffer (0.187 M Tris-HCl pH 6.8, 3 % SDS and 5 mM EDTA) and protein concentrations were determined with a Pierce BCA protein assay kit (Thermo Fisher Scientific, Waltham, MA, USA).

### Cell culture and sustained siRNA treatment with recovery

Caco-2 cells were grown on 10 cm cell culture plates in Dulbecco’s Modified Eagle Medium (DMEM) containing 10 % fetal calf serum, 2 mM L-glutamine, 100 units/ml penicillin and 100 µg/ml streptomycin. The cells were cultured at 37 °C in a 5 % CO_2_ atmosphere. For siRNA experiments, Caco-2 cells were plated on 24- or 12-well plates so that the cells were 20-30 % confluent at the time of siRNA transfection.

siRNAs for human K8 (5’-GCCUCCUUCAUAGACAAGGUA(dTdT)-3’) and K18 (5’-GAGACUGGAGCCAUUACUUCA(dTdT)-3’) and non-target siRNAs based on the K8 siRNA (K8 scramble siRNA; 5′-GUCGUAUAUGACACCGUACCA(dTdT)-3′) and the K18 siRNA (K18 scramble siRNA; 5′-GUAACCAAUGUCGGCGAUACU(dTdT)-3′) were designed in the lab and synthesized (siMAX siRNA, Eurofins Genomics, Ebersberg, Germany). Caco-2 cells were mock-transfected (all reagents except siRNA), transfected with K8 scramble siRNA and K18 scramble siRNA or transfected with K8 siRNA and K18 siRNA using Lipofectamine 2000 (Invitrogen, CA, USA) according to the manufacturer’s instructions (Strnad *et al*., 2016). siRNA transfections were performed using 30 or 60 pmol of each siRNA and 1.5 or 3 μl Lipofectamine 2000 per well in 24- or 12-well plates, respectively. For sustained keratin knockdown, siRNA transfections were repeated three times, followed by 72 h of incubation and subculture after each transfection, and by the end of the third 72-hour incubation samples were collected for protein analysis and immunofluorescence staining. Recovery of cells following siRNA treatment was achieved by incubating previously siRNA-treated cells three times for a 72-hour period which was followed by subculture. Samples were collected for protein analysis at the end of the third incubation period (9 days after the removal of siRNA).

### Immunoprecipitation

Caco-2 cells from two 10 cm cell culture plates were washed with PBS and harvested by scraping. Next, the cells were lysed in 1.5 ml lysis buffer (25 mM Hepes (pH 8.0), 100 mM NaCl, 5 mM EDTA, 0.5% Triton X-100, 20 mM β-glycerophosphate, 20 mM para-nitro-phenyl phosphate, 100 μM ortovanadate, 0.5 mM phenylmethylsulfonyl fluoride, 1 mM dithiothreitol and complete mini protease inhibitor cocktail (Roche, Switzerland)), homogenized by passing the cells through a 23 G needle 10 times and rested 45 minutes on ice. For immunoprecipitation, cell lysates were incubated with rabbit anti-lamin A (Abcam, UK) or mouse anti-K18 (L2A1; Professor Bishr Omary, Rutgers University) antibody (not added to control samples) under rotation at 4 °C overnight. 30 μl of protein-A/G magnetic beads (Thermo Fisher Scientific, Waltham, MA, USA) were added to each sample and the samples were incubated under rotation at 4 °C for 20 minutes. The samples were washed three times in TEG buffer (20 mM Tris-HCl (pH 7.5), 1 mM EDTA, 10% Glycerol, 0.5 mM phenylmethylsulfonyl fluoride, 1 mM dithiothreitol and 1x complete mini protease cocktail (Roche, Switzerland)), dissolved in 3x Laemmli sample buffer and analyzed by SDS-PAGE and Western blot.

### SDS-PAGE and Western blot

Protein samples were homogenized on ice in homogenization buffer (0.187 M Tris-HCl pH 6.8, 3 % SDS and 5 mM EDTA) supplemented with 1x complete protease inhibitor cocktail (Roche, Switzerland) and 1 mM phenylmethylsulfonyl fluoride. Sample protein concentrations were determined with a Pierce BCA protein assay kit (Thermo Fisher Scientific, Waltham, MA, USA) and the samples were normalized and diluted to 5 µg protein/10 µl with 3x Laemmli sample buffer (30 % glycerol, 3 % SDS, 0.1875 M Tris-HCl (pH 6.8), 0,015 % bromophenol blue and 3 % β-mercaptoethanol). The samples were separated on 6-10 % SDS-polyacrylamide gels or 4-20 % Mini-PROTEAN TGX precast gradient gels (Bio-Rad, CA, USA) together with Precision Plus Protein Dual Color Standards (Bio-Rad, CA, USA), transferred to polyvinylidene fluoride membranes and analyzed by Western blot. Primary antibodies used for Western blotting were: rabbit anti-β-actin (Cell Signaling, MA, USA), mouse anti-β-tubulin (Sigma Aldrich, CA, USA), rabbit anti-Emerin (Abcam, UK), rabbit anti-GAPDH (Abcam, UK), rat anti-Hsc70 (Stressgen Bioreagents, MI, USA), rabbit anti-K18 275 (Professor John Eriksson, Finland), rat anti-K19 (Troma III; Developmental Studies Hybridoma Bank, IA, USA) rat anti-K8 (Troma I; Developmental Studies Hybridoma Bank, IA, USA), rabbit anti-Lamin A (Abcam, UK), rabbit anti-Lamin A (Robert Goldman, USA), rabbit anti-phospho Serine 22 - Lamin A/C (Cell Signaling, USA), mouse anti-Lamin A/C (Cell Signaling, MA, USA), rabbit anti-Lamin C (Robert Goldman, USA), mouse anti-Lamin B (Robert Goldman, USA), rabbit anti-Lamin B1 (Abcam, UK), mouse anti-Lamin B2 (Robert Goldman, USA), mouse anti-LAP2α (Roland Foisner, Austria), rabbit anti-Plectin (Gerhard Wiche, Austria), rabbit anti-pRb (Abcam, UK), rabbit anti-phospho-pRb (phospho-Ser 807 and phospho-Ser 811; Cell Signaling, MA, USA), rabbit anti-SUN1 (Abcam, UK), rabbit anti-SUN2 (Abcam, UK) and rabbit anti-YAP (Cell Signaling, MA, USA). Secondary antibodies used for Western blotting were anti-rabbit Alexa Fluor 488/680/800 (Invitrogen, CA, USA), anti-rat Alexa Fluor 488/680 (Invitrogen, CA, USA), anti-mouse Alexa Fluor 488/800 (Invitrogen, CA, USA), anti-rabbit IgG-horseradish peroxidase (HRP) (Promega, WI, USA), anti-rat IgG-HRP (GE Healthcare, UK), anti-rat IgG-HRP (Cell signaling, US) and anti-mouse IgG-HRP (GE Healthcare, UK). HRP-labeled proteins were detected with Amersham ECL Western Blotting Detection Reagent (GE Healthcare, UK) or Western Lightning Plus-ECL (Perkin Elmer, MA, USA) and visualized either on SUPER RX X-ray films (Fuji Corporation, Tokyo, Japan) or using an iBright FL1000 imaging system (Invitrogen, CA, USA). Fluorescently labeled proteins were detected using an iBright FL1000 imaging system (Invitrogen, CA, USA). The Western blot results were quantified using ImageJ software (National Institutes of Health, MD, USA) as previously described (Schneider *et al*., 2012) and normalized to loading controls (Hsc70 or β-tubulin).

### RNA isolation and quantitative RT-PCR

Total lysates of colon tissue were obtained by collecting and combining pieces of PC and DC from 2–5 months old K8^+/+^ and K8^−/−^ male and female mice. The samples were homogenized with a TissueRuptor homogenizer (Qiagen, Germany) and RNA was isolated using a Nucleospin RNA isolation kit (Macherey-Nagel, Germany). Agarose gel analysis was used to assess RNA quality. cDNA was synthesized from 1 μg of each RNA sample using Oligo(dT)_15_ Primer (Promega) and M-MLV Reverse Transcriptase (RNase H Minus, Point Mutant; Promega). Quantitative (q)RT-PCR reactions for target genes were prepared using specific primer (Oligomer, Helsinki, Finland) and probe (Universal probe library, Roche, Basel, Switzerland) combinations (Supplemental Table S1). The target genes were amplified and detected with an Applied Biosystems QuantStudio 3 Real-Time PCR System (Thermo Fisher Scientific) and the gene expression levels were normalized to *β-Actin*. Each cDNA was amplified in triplicates.

### Cryosectioning, fixation and immunofluorescence staining

Pieces of PC and DC from 2–5 months old K8^+/+^ and K8^−/−^ male and female mice were embedded in Tissue-Tek O.C.T compound (Sakura Finetek Europe B.V., Alphen aan den Rijn, The Netherlands) and frozen. The samples were sectioned into 6 µm thick cross sections with a Leica CM3050 S Research Cryostat (Leica Microsystems, Wetzlar, Germany), mounted on microscope slides and fixed with 1 % PFA in PBS, pH 7.4 (Sigma Aldrich, CA, USA) at RT for 10 minutes. Caco-2 cells were cultured on cover slips and fixed similarily with 1 % PFA at RT for 10 minutes. The fixed tissue sections and cell samples were stained as previously described (Ku et al., 2004). The primary antibodies used for immunofluorescence staining were rabbit anti-Lamin A (Abcam, UK), mouse anti-Lamin A/C (Cell Signaling, MA, USA), rabbit anti-Lamin B1 (Abcam, UK), rabbit anti-Myeloperoxidase (MPO; Thermo Fisher Scientific), rat anti-K8 (Troma I, Developmental Hybridoma Bank, NIH, MD; USA), and rabbit anti-YAP (Cell Signaling, MA, USA). The fluorescent secondary antibodies used were anti-mouse Alexa Fluor 488, anti-rabbit Alexa Fluor 488, anti-rat Alexa Fluor 568, anti-rabbit Alexa Fluor 647 (Invitrogen, CA, USA). Draq5 (Cell Signaling, MA, USA) or DAPI (Invitrogen, CA, USA) was used as a nuclear marker. The samples were mounted with ProLong Gold Antifade (Thermo Fisher Scientific). Samples were visualized and imaged at room temperature using a Leica TCS SP5 Matrix confocal microscope (Leica Microsystems) equipped with a 63x Leica PL Apochromat/1.32 oil objective (Leica Microsystems) or a 100x Leica PL Fluotar/1.3 oil objective (Leica Microsystems) using Leica LAS software or a 3i Marianas Spinning disk confocal microscope (Intelligent Imaging Innovations) equipped with a Hamamatsu sCMOS Orca Flash4 v2 C11440-22CU camera (Hamamatsu Photonics, Japan) and a 20x Zeiss Plan-Apochromat/0.8 dry objective (Carl Zeiss AG, Germany) or a 63x Zeiss Plan-Apochromat/1.4 oil objective (Carl Zeiss AG) using Slidebook 6 software. Identical microscopy settings were applied to all samples within one imaging experiment. Images were processed using ImageJ/Fiji and Adobe Photoshop (Adobe, CA, USA) software (Schindelin *et al*., 2012; Schneider *et al*., 2012).

### Senescence-associated β-galactosidase staining

Senescent cells in mouse colon were detected using a Senescence β-Galactosidase Staining Kit (Cell Signaling Technology). Briefly, 10-11.5 months old female K8^+/+^ and K8^−/−^ mice were sacrificed, and PC and DC samples were collected, embedded in Tissue-Tek O.C.T compound and frozen. 4 µm thick sections of PC and DC were produced using a Leica CM3050 S Research Cryostat (Leica Microsystems) and mounted onto microscope slides. The samples were fixed, washed twice with PBS and covered with staining solution. The samples were incubated for 16–18 hours in a dry incubator (no CO_2_), mounted with mowiol and imaged with a Pannoramic Midi FL slide scanner (3DHISTECH, Budapest, Hungary). ImageJ software was used to quantify the percentage of senescent cells by dividing the area of senescence-associated β-galactosidase-positive cells with the total area of the tissue sections.

### Statistical analysis

All quantified Western blot results were statistically analyzed using one way-ANOVA and t-test in GraphPad Prism 5 (GraphPad Software, CA, USA). The results show the mean ± standard deviation (SD) with significant differences shown as * = p < 0.05, ** = p < 0.01 or *** = p < 0.001.

## Acknowledgments

We thank Prof. Gerhard Wiche (Max Perutz Labs, Medical University of Vienna) for the kind donation of the plectin antibody, Assistant professor Pekka Taimen (University of Turku) for the ppRb antibodies and fruitful discussions, Assistant professor Keijo Viiri (University of Tampere) for the Cre-villin mouse strains, and Petra Fichtinger for help with the LAP2α mice (Max Perutz Labs, Medical University of Vienna). We are grateful to all the other members of the Toivola laboratory, especially Frank Weckström, Theresia Jansson, Molly Feiring and Taina Heikkilä (Biosciences/Cell Biology, Faculty of Science and Engineering, Åbo Akademi University; ÅAU), and to members of John Eriksson’s laboratory, especially Elin Torvaldson and Josef Gullmets (Turku Bioscience Centre, University of Turku and ÅAU, and Biosciences/Cell Biology, ÅAU) for fruitful discussions. Imaging was performed at the Cell Imaging and Cytometry Core at Turku Bioscience Centre (University of Turku and ÅAU) and Biocenter Finland. This work was financed by the Academy of Finland 140759/126161 (DMT), Sigrid Juselius Foundation (DMT), Turku Doctoral Programme in Molecular Biosciences at ÅAU (CGAS, JHN), Medicinska Understödsföreningen Liv och Hälsa Foundation (DMT, JHN, CGAS), EU FP7 IRG (DMT), ÅAU Center of Excellence of Cell Stress and Molecular Aging (DMT), The Swedish Cultural Foundation in Finland (JHN, CGAS), Agneta och Carl-Erik Olins foundation (JHN, CGAS), Victoria foundation (CGAS, JHN), K. Albin Johansson foundation (CGAS, JHN), Kommersrådet Otto A. Malms Donationsfond (JHN) and Waldemar von Frenckells foundation (JHN), the EuroCellNet COST Action (CA15214) (DMT, RF), NIH PO1 GM096971 (RDG) and NIH RO106023 (RDG).

## Author contributions

CGAS, JHN, KMR, CBHM, SAA, RDG and DMT conceived and/or designed the experiments; CGAS, JHN, CBH, KR, RF performed the experiments; CGAS, JHN, CBH and DMT analyzed the data; CGAS, JHN and DMT composed the manuscript; CGAS, JHN, DMT, CBH, RF edited the manuscript; CGAS and JHN contributed equally to this manuscript.

## Additional information

## Supplemental Figures and Legends

**Supplemental figure 1.**
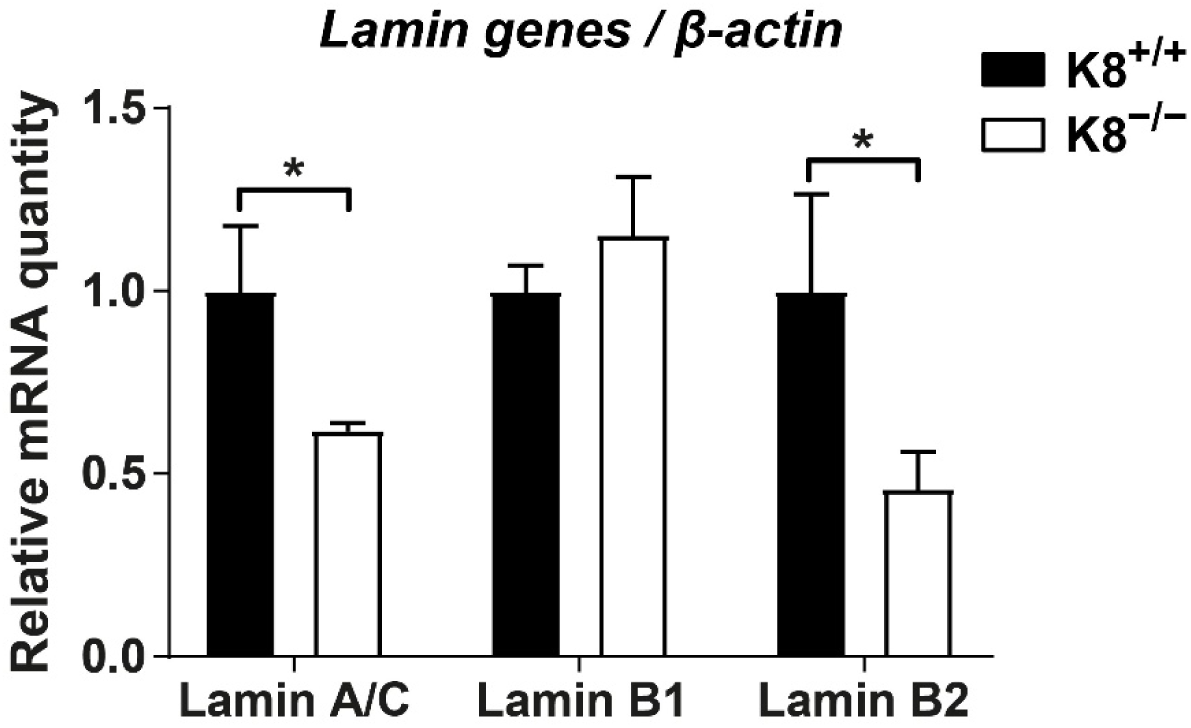
Lamin A/C and B2 mRNA levels are decreased in K8^−/−^ mouse colon. The mRNA levels of lamins A/C, B1 and B2 in K8^+/+^ and K8^−/−^ mouse colon total lysates were analyzed by qRT-PCR. The results were normalized to *β-actin* and represent the average (n=3) fold change ± SD, with significant differences shown as * = p < 0.05.

**Supplemental figure 2.**
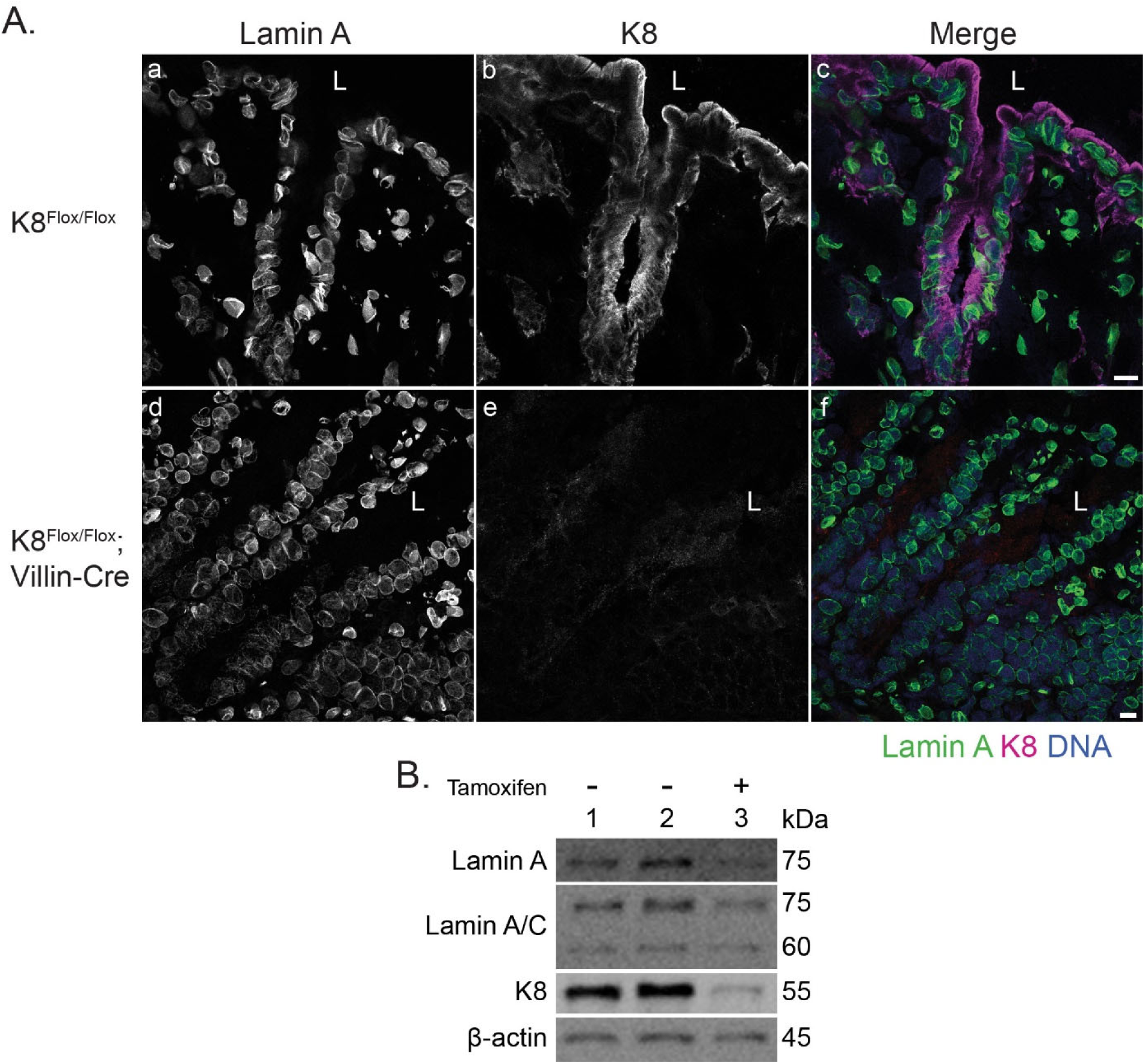
Intestinal epithelial cell-specific knockdown of K8 in mice exhibits colon epithelial cell-specific lamin A downregulation. A) K8^flox/flox^ and K8^flox/flox^;Villin-Cre mouse colon tissue cryosections were immunostained for lamin A (green), K8 (magenta) and DNA (DRAQ5, blue). Scale bar = 10 µm. B) Colon organoids from K8^flox/flox^;Villin-Cre-ER^T2^ mice were isolated, grown in 3D in matrigel, and treated with (+) or without (-) tamoxifen to induce K8 knockdown *ex vivo*. Lysates of the organoids were immunoblotted for lamins A and A/C and K8. β-actin was used as a loading control.

**Supplemental figure 3.**
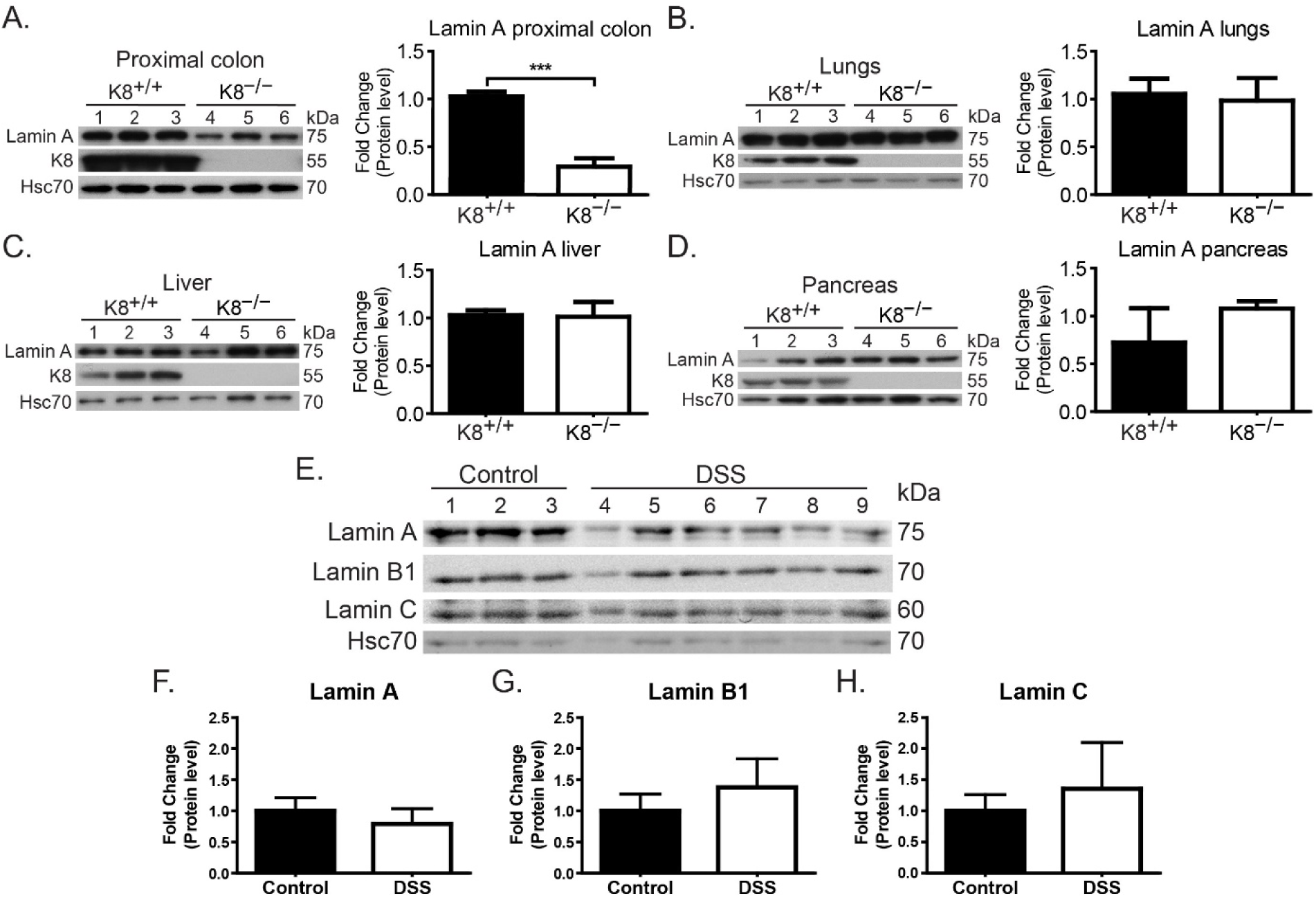
Lamin downregulation is colon-specific in K8^−/−^ mice and not caused by inflammation. Total lysates of proximal colon, lung, liver and pancreas from K8^+/+^ (lanes 1-3) and K8^−/−^ (lanes 4-6) mice (n=3) were immunoblotted for lamin A and K8. Hsc70 was used as a loading control. The immunoblots were quantified and normalized to Hsc70 protein levels and represent the mean (n=3) protein quantity ± SD with significant differences shown as *** = p < 0.001. E-H) Total lysates of colon from untreated (control; lanes 1-3) and DSS-treated (2%, 8 days) K8^+/+^ (lanes 4-9) mice were immunoblotted for lamins A, B1 and C. Hsc70 was used as a loading control. The immunoblots were quantified and normalized to Hsc70 protein levels and represent the mean (n=3) protein quantity ± SD.

**Supplemental figure 4.**
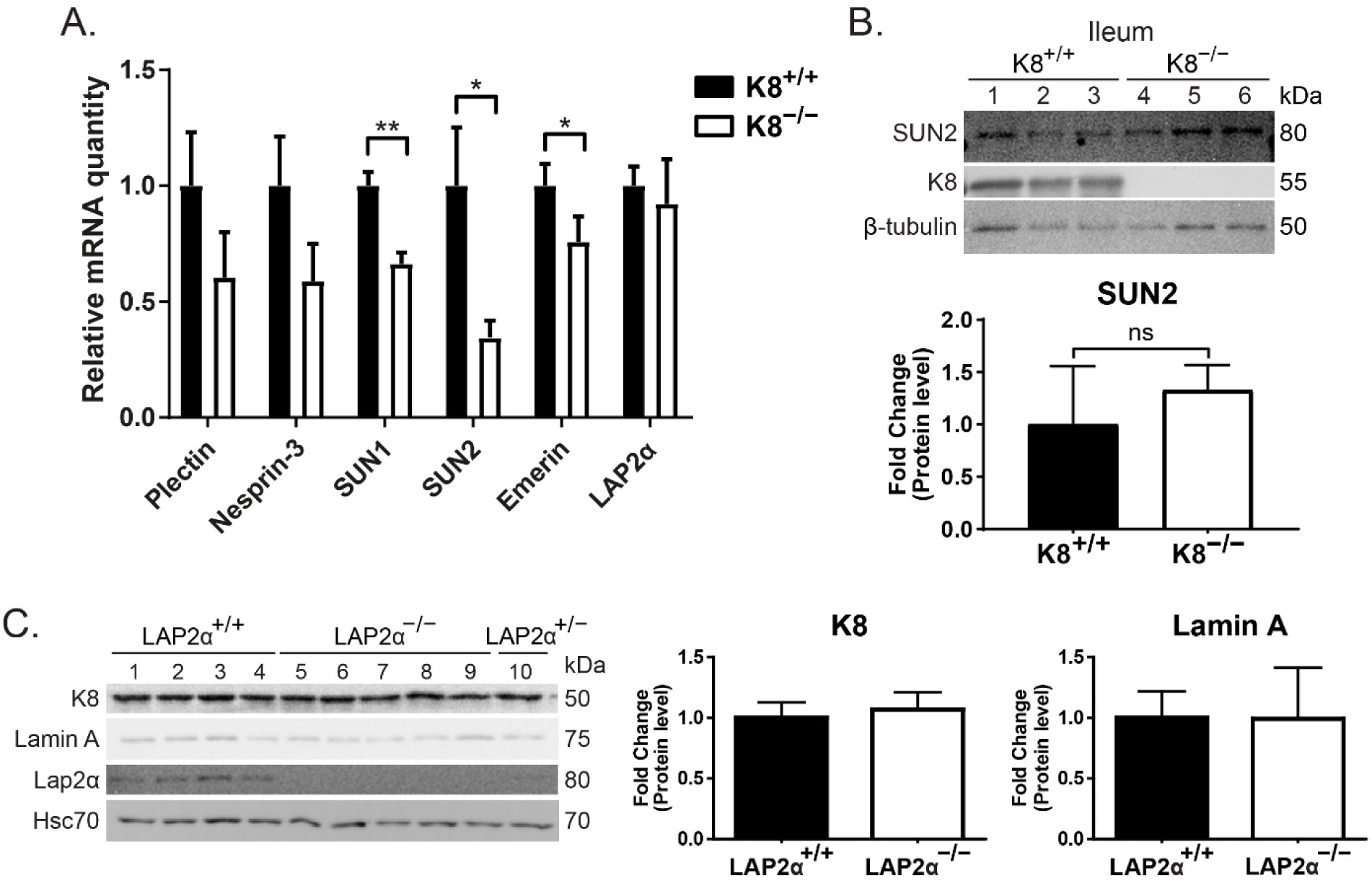
LINC protein down-regulation is colon-specific in K8^−/−^ mice and partly due to reduced gene expression, while keratins and lamin A are unaltered in LAP2α^−/−^ mouse colon. A) The mRNA levels of plectin, nesprin-3, SUN1, SUN2, emerin and LAP2α in K8^+/+^ and K8^−/−^ mouse colon total lysates were analyzed by qRT-PCR. The results were normalized to *β-actin* and represent the average (n=3) fold change ± SD, with significant differences shown as * = p < 0.05 and ** = p < 0.01. B) Total lysates of ileum from K8^+/+^ (lanes 1-3) and K8^−/−^ (lanes 4-6) mice were immunoblotted for SUN2 and K8. β-tubulin was used as a loading control. The immunoblots were quantified and normalized to β-tubulin protein levels and represent the mean (n=3) protein quantity ± SD. C) Lysates of crudely isolated colon epithelium from LAP2α^+/+^ (lanes 1-4), LAP2α^−/−^ (lanes 5-9) and LAP2α^+/–^ (lane 10) mice were immunoblotted for K8, lamin A and LAP2α. Hsc70 was used as a loading control. The immunoblots were quantified and normalized to Hsc70 protein levels and represent the mean (n=4 for LAP2α^+/+^ and n=5 for LAP2α^−/−^) protein quantity ± SD.

**Supplemental figure 5.**
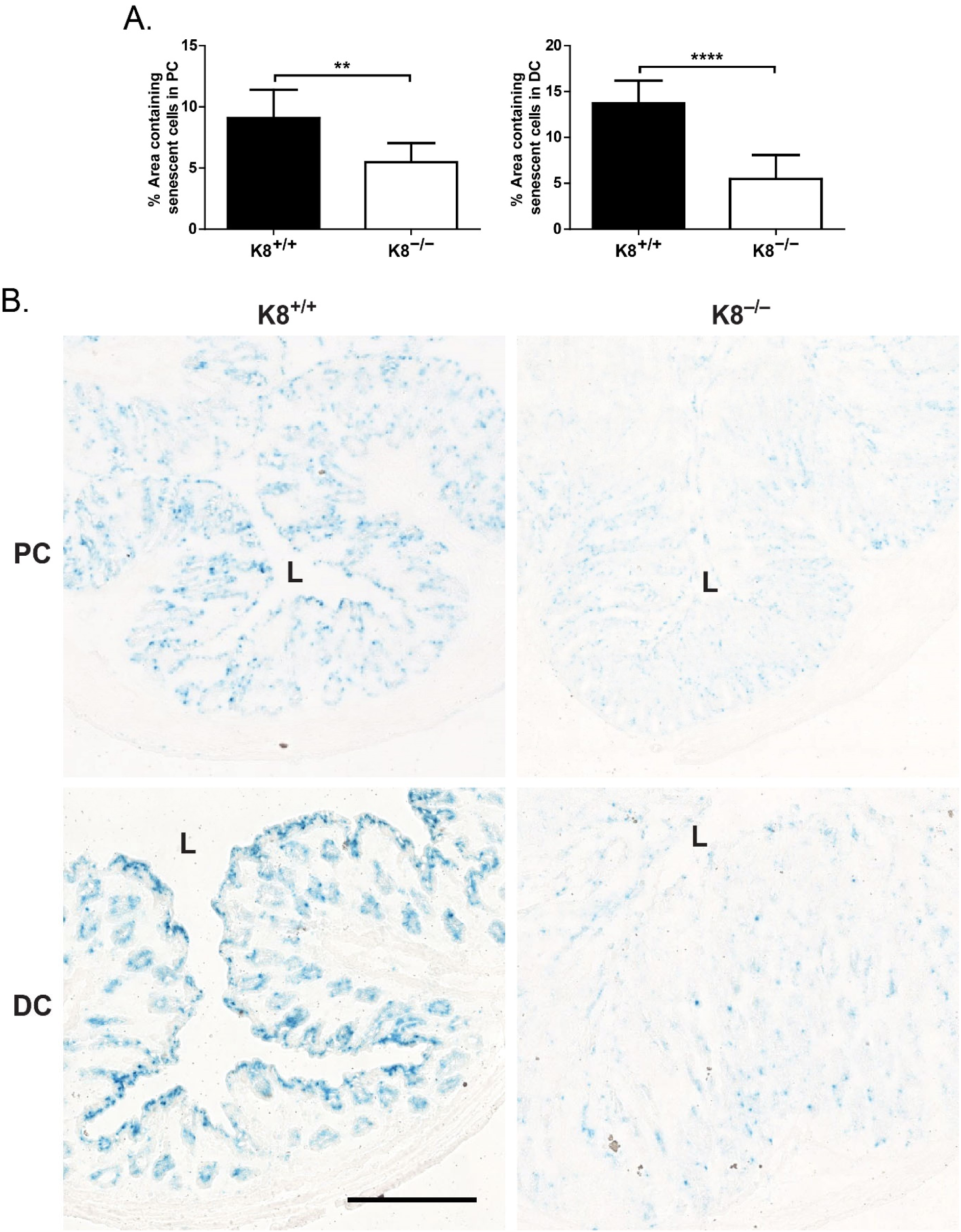
Cell senescence is diminished in the K8^−/−^ mouse colon. Tissue sections of proximal (PC) and distal (DC) colon from K8^+/+^ and K8^−/−^ mice were analyzed for the presence of senescent cells by senescence-associated β-galactosidase staining. A) The number of senescent cells was quantified with ImageJ software by dividing the area of senescence-associated β-galactosidase-positive blue signal with the total area of the tissue section. B) Representative images of K8^+/+^ and K8^−/−^ PC and DC with senescent cells in blue. Scale bar = 500 µm. L= lumen.

**Supplemental table 1.**
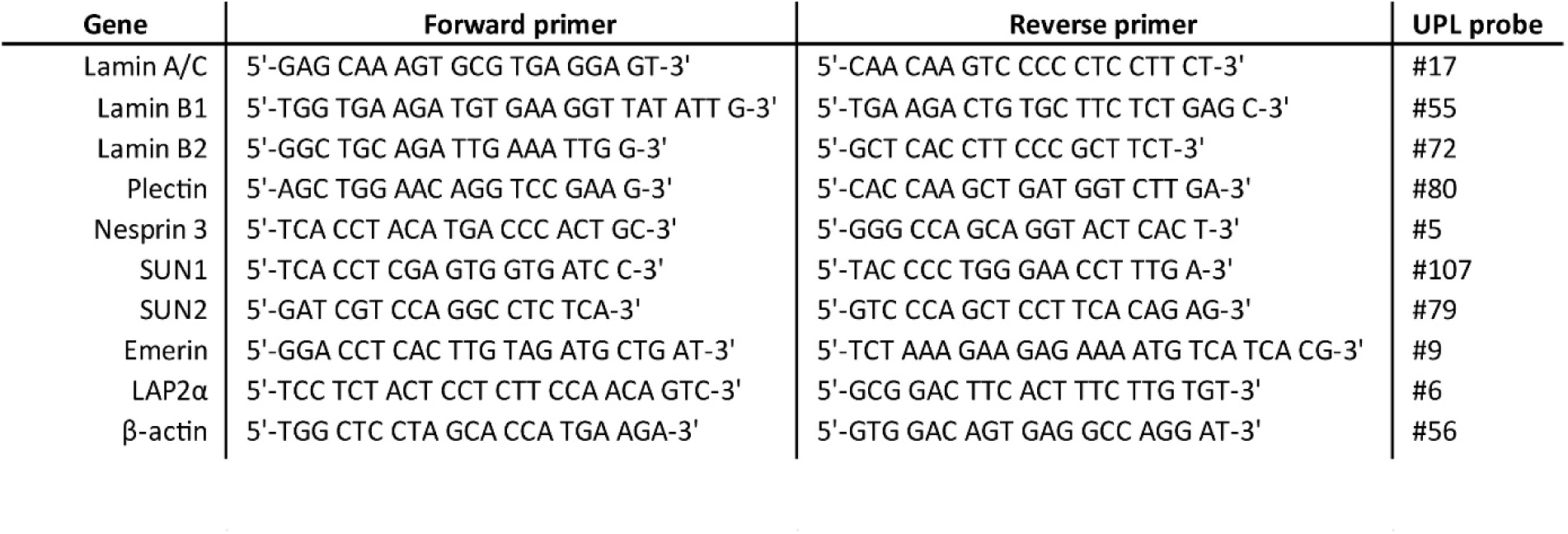
Primers and Universal ProbeLibrary (UPL) probes used for quantitative RT-PCR.

